# Requirements for establishment and epigenetic stability of mammalian heterochromatin

**DOI:** 10.1101/2023.02.27.530221

**Authors:** Antonis Tatarakis, Harleen Saini, Danesh Moazed

## Abstract

Heterochromatic domains of DNA account for a large fraction of mammalian genomes and play critical roles in silencing transposons and genes, but the mechanisms that establish and maintain these domains are not fully understood. Here we use an inducible heterochromatin formation system combined with a CRISPR-based genetic screen to investigate the requirements for the establishment and maintenance of heterochromatin in mouse embryonic stem cells (mESCs). We show that DNA sequence-independent and histone H3 lysine 9 methylation (H3K9me)-dependent heterochromatin can be inherited for a limited number of cell divisions in mESCs but becomes stable upon differentiation. We provide evidence that the increased stability of heterochromatin in differentiated cells results from the downregulation of one or more enzymes that erase H3K9me and DNA methylation. Moreover, we show that in addition to components of the H3K9 and DNA methylation pathways, heterochromatin maintenance requires DHX9 and other RNA processing proteins. DHX9 is an RNA/DNA helicase with previously described roles in preventing genomic instability resulting from transcription-associated replication stress. We found that deletion of DHX9 results in defective heterochromatin inheritance and is associated with increased transcription of major satellite repeats, accumulation of R-loops, and loss of H3K9me. Our findings define the requirements for the establishment and epigenetic inheritance of mammalian heterochromatin and suggest that R-loops and replication stress lead to epigenetic instability.

## Introduction

Heterochromatin is a conserved feature of eukaryotic chromosomes that comprises nearly half of the genome in some eukaryotes and has important functions in regulation of transcription, silencing of repetitive DNA sequences, and maintenance of genome integrity^1-4^. Establishment of heterochromatic domains is initiated by the DNA sequence-dependent recruitment of histone-modifying enzymes to nucleation sites, followed by spreading of the modification to nearby regions by a read-write mechanism involving iterative nucleosome binding and modification cycles^3, 5, 6^. Studies in fission yeast have uncovered roles for specific DNA sequences in epigenetic inheritance of histone H3 lysine 9 methylation (H3K9me)-associated heterochromatin^7–10^. Whether H3K9me can mediate epigenetic inheritance in mammalian cells in the absence of other inputs remains unknown.

In mammalian cells, H3K9me is required for the formation of constitutive heterochromatin at pericentromeric regions, telomeric repeats, and ribosomal DNA repeats^11, 12^. Six methyltransferase enzymes are responsible for H3K9me catalysis in mammalian cells with distinct modes of action. Although the complex interplay between H3K9 methyltransferases is not well understood, deletion of all six enzymes is required to completely disrupt heterochromatin organization highlighting their unique roles^13^. Following its establishment, H3K9me provides binding sites for the highly conserved heterochromatin protein 1 (HP1) family members, which serve as a platform for recruitment of downstream effector proteins^14–16^. Additional epigenetic modifications have been linked to heterochromatin in mammalian cells, such as DNA 5-methylcytosine (5mC), which colocalizes with H3K9me at heterochromatic domains, and is required for silencing retroviral elements in mammals^17–19^. Whether DNA methylation is required for silencing of H3K9me enriched regions and participates in the maintenance of H3K9me heterochromatin is unknown.

Heterochromatic domains are not transcriptionally inert and give rise to noncoding RNAs, which contribute to the assembly of heterochromatin. Studies in fission yeast, plants and various animals have demonstrated the function of small RNAs and the RNA interference (RNAi) pathway in establishing heterochromatin via RNAi interactions with nascent RNA to promote the recruitment of H3K9me enzymes at specific genomic regions^20–23^. In mammalian cells, nascent repeat-associated RNAs facilitate the recruitment of H3K9me enzymes and the establishment of heterochromatin^24–28^. Such an RNA-based mechanism appears to complement the role of site-specific DNA binding factors in heterochromatin assembly. A recent study in fission yeast demonstrated that degradation of heterochromatin-associated RNAs by the rixosome complex contributes to H3K9me maintenance, further highlighting the role of RNA processing in epigenetic inheritance of heterochromatin^29^. Transcription and noncoding RNAs are also components of mammalian heterochromatin, but the molecular mechanism responsible for their regulation and their role(s) in heterochromatin maintenance are not fully understood.

Previous studies have demonstrated that ectopic domains of H3K9me3 heterochromatin can be established and stably maintained in mammalian cells^30–32^. Here we use an inducible heterochromatin assembly system to investigate the requirements for the establishment and epigenetic inheritance phases of heterochromatin formation. We show that in mouse embryonic stem cells (mESCs), heterochromatin can be inherited independently of the underlying DNA sequence input but only for a limited number of cell divisions and that this metastable mode of inheritance becomes remarkably stable upon differentiation. Furthermore, to comprehensively assess the importance of nuclear factors in the establishment and epigenetic inheritance of heterochromatin, we performed a two-tiered CRISPR-Cas9-based genetic screen. We identified numerous factors required for heterochromatin establishment and those required solely for maintenance. Our findings demonstrate that multiple H3K9 and DNA methylation enzymes cooperate with downstream factors to establish and maintain heterochromatin. Among the maintenance-specific factors, we show that the loss of DHX9, an RNA/DNA helicase required for R-loop resolution^33–35^, greatly diminishes epigenetic inheritance of the inducible heterochromatin domain and impairs the maintenance of H3K9me3 and silencing at major satellite repeats.

## Results

### Inducible system to separate establishment and maintenance of silencing in mESCs

To determine whether mammalian heterochromatin maintenance can be separated from the sequences that initiate its establishment, we developed a system for inducible heterochromatin formation in mESCs. We fused the bacterial tetracycline repressor (TetR) protein to the KRAB (Krüppel-associated box) domain found in many zinc finger proteins, which recruits the histone H3K9 methyltransferase SETDB1 to initiate silencing^36, 37^. The TetR DNA binding domain facilitates the targeting of this fusion protein to a locus that harbors its cognate DNA binding sequence, tetracycline Operator (*tetO*). The addition of doxycycline (+Dox medium) releases the TetR fusion protein from *tetO* sites, allowing us to separate the sequence-dependent phase of heterochromatin establishment from the sequence-independent maintenance phase (Figure 1a). To generate a reporter locus and monitor its silencing in mESCs, we inserted 8 *tetO* sites (*8x-tetO*) immediately upstream of a mammalian promoter driving the expression of enhanced green fluorescence protein (eGFP) (Figure S1a, Figure 1a). This reporter gene was inserted at a euchromatic locus, which contained active transcription-related histone modifications, such as histone 3 lysine 4 trimethylation (H3K4me3) and histone 3 lysine 36 trimethylation (H3K36me3), lacked H3K9me3 (Figure S1a), and expressed mRNA in mESCs and across different tissues (Figure S1a). We reasoned that such a locus would lack sequence elements that may contribute to heterochromatin maintenance or spontaneous silencing, often observed when reporter genes are inserted at gene deserts (data not shown). We further engineered mESCs to express either a Flag-tagged TetR-KRAB domain protein (TetR-Flag-KRAB) or Flag-tagged TetR as a control (TetR-Flag) (Figure S1b). We then tested the effect of the fusion proteins (Figure 1b) on the expression of the *8x-tetO-eGFP* reporter gene by monitoring eGFP^+^ cells by fluorescence-activated cell sorting (FACS).

**Figure 1.**
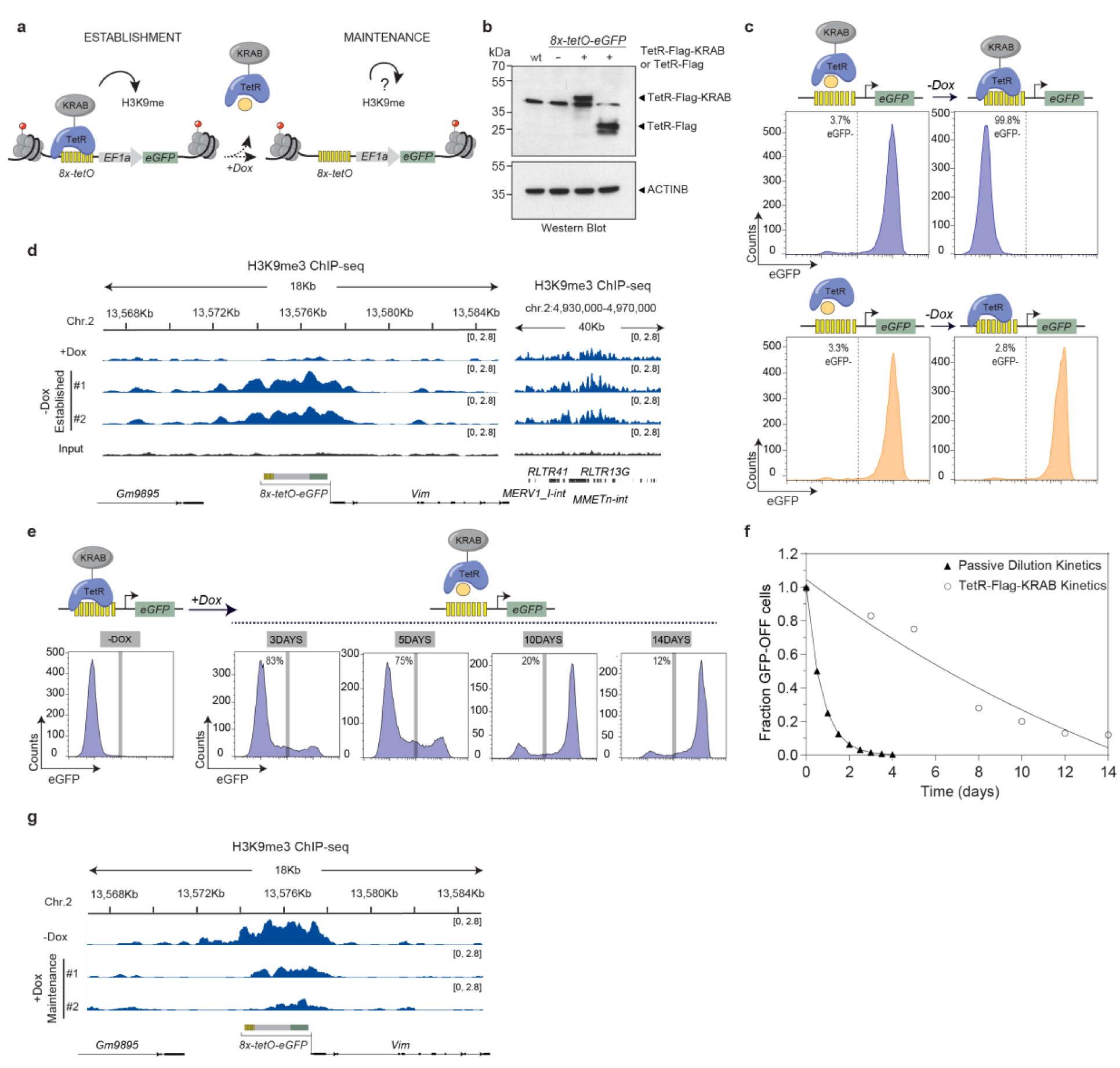
Maintenance of inducible silencing and H3K9me3 in mESCs. **a,** Schematic diagram of the inducible reporter system for heterochromatin establishment and maintenance. EF1a, elongation factor 1a promoter; *8x-tetO*, 8 copies of the *tet* operator; TetR-KRAB, fusion of TetR and KRAB domain; H3K9me, tri-methyl histone H3 Lys 9; DOX, doxycycline. **b,** Western blot showing protein levels of TetR-Flag-KRAB and TetR-Flag (top) in mESC lines generated with the insertion of the respective transgenes at the *Rosa26* locus in the *8x-tetO-eGFP* reporter mESC line and β-Actin (bottom) western blot as a loading control. Molecular weights in kilodalton are shown on the left. **c,** Flow cytometry histograms show eGFP expression before and 10 days after recruitment of TetR-Flag-KRAB (blue) or TetR-Flag (orange) at the reporter locus. Top, schematic diagram shows the experimental design. Percentages (%) indicate fraction of eGFP^-^ cells. **d,** Genome tracks of ChIP-seq for H3K9me3 and input for the region surrounding the reporter locus (yellow, *8x-tetO* sites; grey, EF1a promoter; green, eGFP) before (+Dox) and after establishment (-Dox) of silencing. Normalized reads are presented in brackets. Top, chromosome coordinates; Right, control showing enrichment of H3K9me3 at native ERV elements. **e,** Flow cytometry histograms showing eGFP expression after adding back doxycycline (+Dox) to the growth medium of mESCs with an established silent state at the reporter locus to assess maintenance of silencing at the indicated time points. **f,** Decay kinetics of reporter locus silencing during the maintenance assay. Fraction of GFP-OFF cells at various time points after addition of doxycycline (+Dox) to cells with an established silenced state (o) and simulated data for the theoretical maintenance of the silent state if only passive dilution of the modified histones occurs ( ). Exponential curve fitting was used as a guide. **g,** Genome tracks of ChIP-seq for H3K9me3 for the region surrounding the reporter locus (as in **d**) after establishment (-Dox) of silencing and 5 days after the addition of doxycycline (+Dox) back to the medium to assess maintenance. Normalized reads are presented in brackets. Note the different scale used. Top, chromosome coordinates.

In the absence of Dox (-Dox medium), eGFP reporter was expressed in cells containing TetR or the reporter alone but strongly silenced in cells with TetR-KRAB (TetR-KRAB cells) (Figure 1c, Figure S1c). Chromatin immunoprecipitation (ChIP) combined with high-throughput sequencing (ChIP-seq) or quantitative polymerase chain reaction (ChIP-qPCR) showed the establishment of an 8-to 10-kb H3K9me3 domain spanning the reporter gene region in TetR-KRAB cells cultured in -Dox medium (Figure 1d, Figure S1d). H3K9me3 enrichment at the reporter locus was similar to that observed at the native intracisternal A particle (IAP) and ERV heterochromatic loci (Figure 1d, Figure S1d). H3K4me3 levels followed the opposite trend, decreasing to background levels in TetR-KRAB cells (Figure S1e). To assess whether the eGFP^-^ silent state could be maintained in the absence of continuous establishment, the cell culture medium was supplemented with Dox to release TetR-KRAB from the *tetO* sites. Quantification of eGFP^+^ and eGFP^-^ cells revealed that the silent state persisted in ∼85% of the cells after 3 days of growth in +Dox medium (∼5-6 cell divisions), in ∼75% of the cells after 5 days (∼9-10 cell divisions), in ∼20% of the cells after 10 days (∼18-20 cell divisions), and in ∼15% of cells after 14 days (∼25-28 cell divisions) (Figure 1e, f). The decay rate of the silent state was slower than what would be expected from residual silencing due to passive dilution of histone modifications, indicating an active mechanism of propagation (Figure 1f). Consistent with eGFP expression, ChIP-seq and ChIP-qPCR showed that the H3K9me3 domain spanning the reporter region was maintained after 5 days of growth in +Dox medium but was decreased in size and magnitude (Figure 1g, Figure S1f). This observation is consistent with the loss of silencing in 20% of cells at this time point. Control ChIP experiments showed that TetR-KRAB and TetR were recruited to the *8x-tetO* sites in -Dox medium and released after switching to +Dox medium (Figure S1g). This binding was specific and not detected at a negative control locus (*Gapdh*) (Figure S1h). To rule out the possibility that H3K9me3 maintenance may reflect residual or weak binding of TetR-KRAB to the *8x-tetO* sites, we used CRISPR-Cas9 to delete the TetR-KRAB expression cassette. Deletion of TetR-KRAB prevented silencing of the eGFP reporter gene (Figure S1i, j). However, the eGFP silent state was maintained when we deleted TetR-KRAB via lentiviral-mediated CRISPR-Cas9 genome editing 7 days after establishing a silent state (Figure S1k, l). Furthermore, the fraction of eGFP^-^ cells after Dox addition was similar between wild-type and TetR-KRAB-depleted cells (Figure 1f, Figure S1l), ruling out heterochromatin maintenance due to residual TetR-KRAB binding to the *8x-tetO* sites. These results demonstrate that a newly established H3K9me domain and its associated silent state can be maintained in the absence of sequence-dependent initiation through a limited number of mitotic cell divisions in mESCs.

### A CRISPR screen identifies factors essential for the establishment of silencing

To identify factors important for H3K9me-dependent silencing in mESCs, we performed a forward CRISPR-Cas9 genetic screen using our inducible silencing system. We generated a pooled library of 5950 single guide RNAs (sgRNAs) targeting 1160 genes (5 sgRNAs/gene and 150 non-targeting control sgRNAs; “EpiChromo” Library), including known chromatin and epigenetic regulators, DNA replication factors, nuclear periphery factors, and RNA processing factors. We also included factors identified in proteomic analyses to be associated with heterochromatin but whose role in chromatin regulation and function remains unknown (Figure S2a, Supplementary Tables 1, 2). Using lentiviral transduction, we introduced sgRNAs into our engineered mESCs before establishment of a silent state (eGFP^+^) such that each cell incorporated a single sgRNA (Figure 2a). We then switched to -Dox medium to establish reporter gene silencing (Figure 2a). After 10 days of establishment in -Dox medium, we isolated eGFP^+^ cells by FACS, presuming that each cell contained a mutation in a factor essential for the establishment of silencing. We then performed high-throughput sequencing to assess the frequencies of the individual sgRNAs in the eGFP^+^ cells, as well as in unsorted cells for comparison. We used the Model-based Analysis of Genome-wide CRISPR-Cas9 Knockout (MAGeCK) robust ranking aggregation (RRA) algorithm for hit identification and statistical analysis of the results (Figure 2a)^38^.

**Figure 2.**
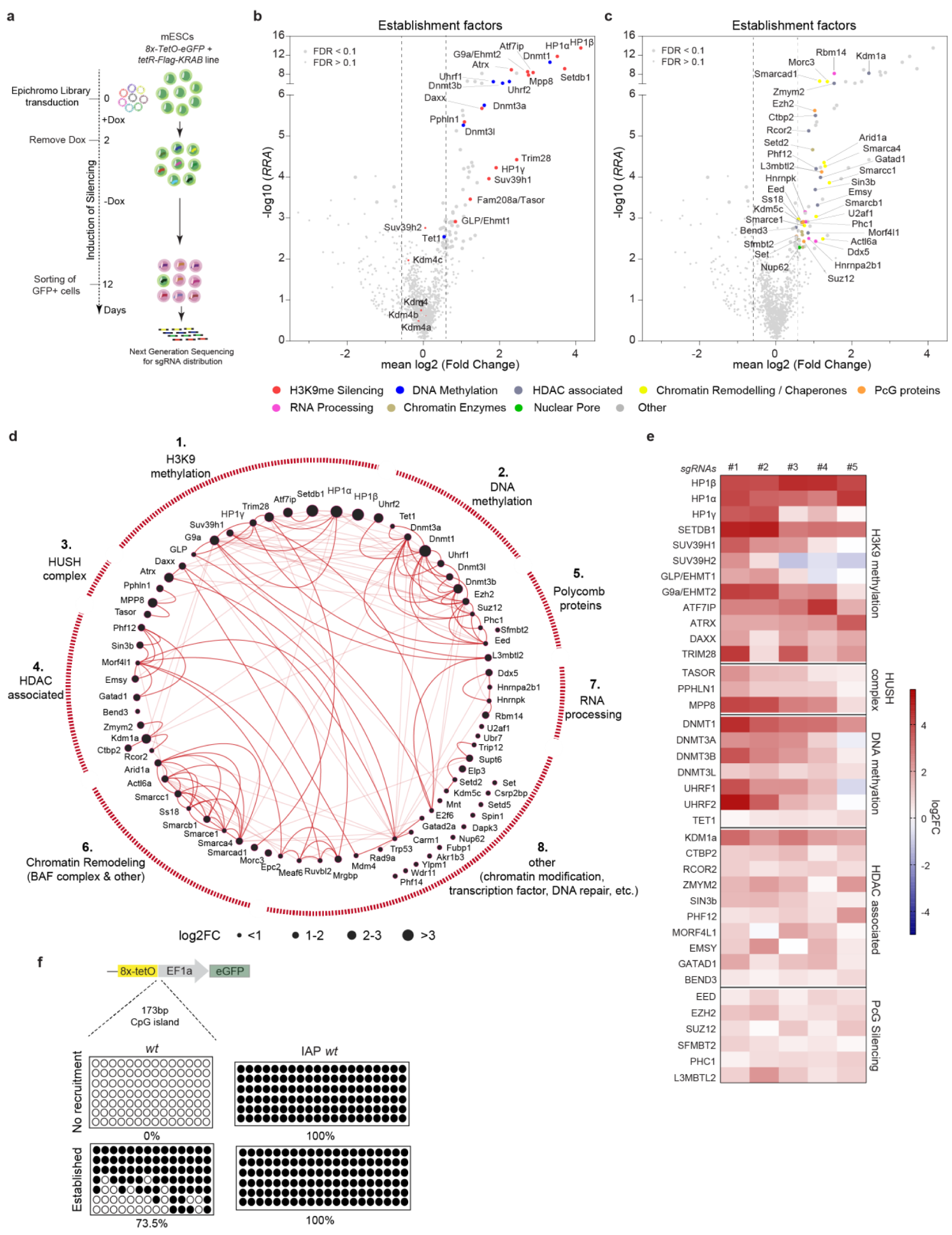
A CRISPR screen identifies mutants deficient for silencing establishment. **a,** Schematic of the pooled screen strategy for identification of genes required for the establishment of silencing. Mouse ES cells carrying the *8x-tetO-eGFP* reporter gene and the TetR-Flag-KRAB inserted at the *Rosa26* locus were transduced with the EpiChromo viral pooled library. Doxycycline was removed (-Dox) from the medium to induce silencing of the reporter gene. At the end of the establishment period, eGFP^+^ cells were isolated and the sgRNA sequences of sorted and unsorted cells were sequenced to quantify the abundance of individual sgRNAs. **b,** Volcano plot displaying the screen results for establishment of silencing. The plot shows the mean values of log_2_ fold ratios of all 5 sgRNAs targeting each gene in eGFP^+^ sorted versus unsorted cells (x axis) plotted against the negative log_10_ significant score (RRA) values (y axis) (n = two replicates). Dashed vertical black lines mark the mean log_2_ (Fold Change) -0.585 (left) and 0.585 (right). Selected chromatin-associated proteins are highlighted and grouped into known functional categories (color code under the plots). Source data for this figure are provided in Supplementary Table 3. **c,** Same as in (**b**) showing additional functional groups of proteins affecting the establishment of silencing. **d,** STRING clustering of 79 factors identified to affect establishment of silencing at the reporter locus (FDR < 0.1). Functional interactions based on STRING database are shown with red lines between proteins. The darker red lines depict interactions between clustered proteins uncovered by Markov clustering (MCL). **e,** Heatmap of gene hits (left) affecting the establishment of silencing belonging to the indicated functional categories (right). Log_2_ fold change of each one of the 5 sgRNAs targeting each gene in eGFP^+^ sorted versus unsorted cells is shown (n = two independent replicates). **f,** Bisulfite sequencing results. Top, schematic diagram of the reporter gene. Bottom, bisulfite sequencing of DNA CpG methylation at the reporter locus before recruitment of the TetR-Flag-KRAB and after establishment of the silent state. Filled and open circles represent methylated and unmethylated CpGs, respectively. Intra-cisternal A-type (IAP) sequences were used as internal control. Percentages (%) indicate methylated CpGs.

We identified 79 genes required for the establishment of eGFP silencing, at a 10% false discovery rate (FDR) cut-off requiring at least 3 of 5 sgRNAs per target enriched in the eGFP^+^ pool (Figure 2b, c; Supplementary Table 3). Gene ontology analysis revealed an enrichment of genes with known functions in heterochromatin formation, DNA methylation, and histone methylation, as well as broader transcriptional regulation and chromatin remodeling functions (Figure 2b, c; Figure S2b, c). We generated lentiviral-mediated knockout mESC lines to validate the screen results using different sgRNAs for 17 selected candidate factors. All 17 mESC lines showed increased expression of the eGFP reporter relative to mESCs generated using control sgRNAs (Figure S2d).

The factors required for heterochromatin establishment were grouped into 8 clusters based on functional interactions in the STRING database (Figure 2d). Cluster 1 included proteins associated with H3K9 methylation, HP1 proteins (HP1α, HP1β, HP1γ), H3K9 methylation enzymes (SETDB1, G9a/EHMT2, GLP/EHMT1, SUV39H1), SETDB1 interaction partners (ATF7IP and KRAB-associated protein 1, TRIM28/KAP1), and the ATRX-DAXX histone chaperone complex (Figure 2b-e, Figure S2e). Notably, *Suv39h2* mutations showed weaker defects in silencing (FDR = 0.18) with 2 out of 5 sgRNAs being significantly enriched in eGFP^+^ cells (Figure 2e). Cluster 2 included DNA methyltransferases (DNMT1, DNMT3A, DNMT3B, DNMT3L), UHRF1, and 2 other proteins. Consistent with the requirement of DNA methylation enzymes and co-factors in the establishment of silencing, 5-methylcytosine (5mC) bisulfite sequencing at the promoter of the reporter gene showed that nearly 75% of CpG dinucleotides were methylated (Figure 2f). Cluster 3 included components of the HUSH complex (FAM208A/TASOR, PPHLN1, MPP8) (Figure 2b-e, Figure S2e). Cluster 4 contained proteins associated with histone deacetylation enzymes (HDACs). We identified components of the SIN3/HDAC complex such as SIN3b, MORF4L1, PHF12, EMSY, and GATAD1, as well as components of the coREST/HDAC complex such as RCOR2, KDM1A/LSD1, ZMYM2, and CTBP2 (Figure 2c-e, Figure S2e, Supplementary Table 3). Notably, HDACs represented in the EpiChromo library were not uncovered as positive hits in our screen (Figure S2f). Heterochromatic domains are generally hypoacetylated, and the role of HDACs and their complexes in heterochromatin formation and silencing is well documented^23, 39^. Therefore, our results most likely reflect redundancy between the many different HDAC enzymes in mESCs. These results suggest that the mechanism of mammalian H3K9me heterochromatin assembly is highly cooperative and requires input from multiple histone H3K9 enzymes, DNA methyltransferases, and HDACs complexes which can only partially compensate for each other.

Cluster 5 included components of complexes associated with Polycomb-dependent gene silencing (SUZ12, EZH2, EED, PHC1, SFMBT2, L3MBTL2), although their effects were not as robust as those of the core H3K9me and DNA methylation proteins (Figure 2c-e, Figure S2e)^40^. A functional interplay between H3K9me and DNA methylation with Polycomb-mediated silencing has been previously reported^41–44^. To test whether the reporter locus gains H3K27me3 after establishment of silencing, we performed H3K27me3 ChIP-qPCR. We found increased enrichment, although at a lower magnitude relative to a native genomic locus associated with H3K27me3 (Figure S2g). These results suggest that at least in some cells, or to some extent, H3K9me3 and H3K27me3 pathways may cooperate to silence the reporter gene. Alternatively, the weaker silencing of the reporter locus in cells carrying mutations of Polycomb proteins may be explained by the redistribution of the H3K9me machinery from the reporter locus to H3K27me genomic targets.

Cluster 6 contained several BAF complex components (SMARCA4, SMARCB1, SMARCE1, ACTl6A, ARID1A, SMARCC1, SS18) and other chromatin remodelers including the Swi/Snf-related protein SMARCAD1 and the MORC family CW-type zinc finger protein MORC3 (Figure 2c-e, Figure S2h), some of which have been previously shown to play a role in retroviral silencing^45–48^. Cluster 7 included several proteins associated with pre-mRNA processing, suggesting interactions between nascent RNA regulatory pathways and heterochromatin-mediated gene silencing. For example, DDX5 (RNA helicase involved in pre-mRNA splicing, which also acts as a corepressor together with HDAC1)^49^, RBM14 (RNA binding protein regulator of splicing and transcriptional corepressor), HNRNPK and HNRNPA2B1 (pre-mRNA binding proteins), and U2AF1 (U2 snRNA auxiliary factor 1) were all required for reporter gene silencing (Figure 2c-d, Figure S2i). Cluster 8 represented proteins that may affect heterochromatin directly or indirectly, such as KDM5C, which similarly to KDM1A/LSD1 (cluster 4) histone demethylase, removes methyl marks associated with transcription, as well as the SET protein, which as a component of the inhibitor of acetyltransferase activity (INHAT) complex has been proposed to inhibit histone acetylation and may be required for efficient heterochromatin assembly (Figure 2c-d, Figure S2j)^50–52^. Other proteins (e.g., E2F6, TRP53, MDM4) in this cluster have been functionally associated with proteins involved in the H3K9 methylation pathway (Figure 2d). Therefore, in addition to input from H3K9 and DNA methyltransferases, the HUSH complex, and Polycomb proteins, establishment of an H3K9me3-dependent domain and silencing require chromatin remodeling and pre-mRNA processing.

### Identification of genes required for epigenetic inheritance of silencing

We next performed a CRISPR-Cas9 screen for factors required for the sequence-independent maintenance phase of silencing. After transducing mESCs carrying the inducible eGFP reporter with the EpiChromo library, we allowed establishment of silencing for 10 days and selected eGFP^-^ cells (silenced) by FACS (Figure 3a). To screen for maintenance factors, eGFP^-^ cells were then switched to +Dox medium (to release TetR-KRAB from the *8x-tetO* sites) for 3 days. eGFP^+^ cells were selected by FACS and the enrichment of individual sgRNAs was evaluated as described above for the establishment screen. Although there is some spontaneous decay of silencing and H3K9me3 at this time point, the fraction of wild-type cells that lose the silent state is low (<20%) and minimally affects the sensitivity of our assay (Figure 1e, f).

**Figure 3.**
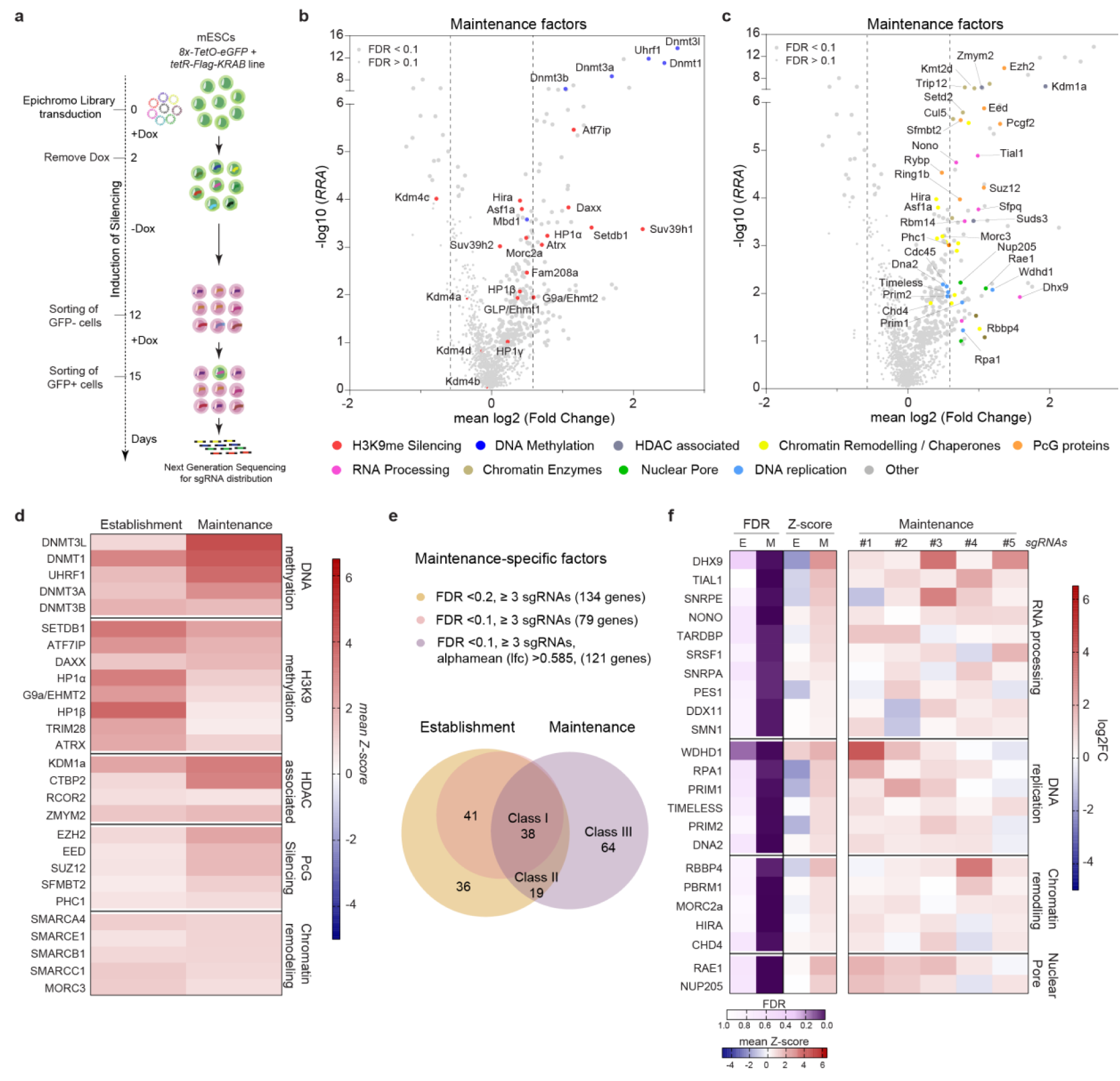
Genes required for epigenetic inheritance of silencing. **a,** Schematic of the pooled screen strategy for identification of genes required for maintenance of silencing. Mouse ES cells carrying the inducible heterochromatin system were transduced with the EpiChromo viral pooled library and following establishment of silencing eGFP^-^ cells were sorted out. Doxycycline was added back to the medium (+Dox) to release TetR-KRAB from the *8x-tetO* sites and after 3 days, eGFP^+^ cells were sorted out by FACS to quantify the abundance of individual sgRNAs by sequencing. **b,** Volcano plot displaying the screen results for the maintenance of silencing plotted as in Figure 2b. Selected chromatin-associated proteins are highlighted and grouped into known functional categories (color code under the plots). Source data for this figure are provided in Supplementary Table 4. **c,** Same as in **b** showing additional functional groups of proteins affecting maintenance of silencing. **d,** Heatmap of gene hits (left) belonging to the indicated functional categories (right) that affect both establishment and maintenance of silencing. Mean values of the Z-scores of all 5 sgRNAs targeting each gene in eGFP^+^ sorted versus unsorted cells is shown in each condition (n = two independent replicates). **e,** Venn diagram showing the number of genes affecting the establishment, maintenance, or both phases of silencing of the reporter locus. Alphamean, mean of log_2_ fold change of the effective sgRNAs. **f,** Heatmaps of gene hits (left) belonging to the indicated functional categories (right) that are maintenance-specific regulators of silencing. Left, gene-level FDR values and mean values of the Z-scores of all 5 sgRNAs targeting each gene in eGFP^+^ sorted versus unsorted cells during establishment (E) or maintenance (M) is shown. Right, log_2_ fold change of each one of the 5 sgRNAs targeting each gene in eGFP^+^ sorted versus unsorted cells is shown (n = two independent replicates).

This screening strategy identified 121 candidate genes required for maintenance of the silent state using a 10% FDR as a cut-off and requiring at least 3 of 5 sgRNAs per target enriched in the eGFP^+^ cells (with an average log_2_ fold change >0.585 for eGFP^+^ cells versus unselected cells) (Figure 3b, c, Supplementary Table 4). We further grouped these genes as class I (38 hits, required for both establishment and maintenance), class II (19 hits, maintenance factors with weak effects on establishment), and class III (64 hits, maintenance-specific). Class I factors included DNMT1, UHRF1, DNMT3A, DNMT3L, DNMT3B, and H3K9me-associated proteins, SETDB1, ATF7IP, the ATRX-DAXX complex, HP1α, HP1β, G9a/EHMT2, and TRIM28 (Figure 3b, d, and Figure S3c). Overall, DNMT mutations had the strongest maintenance defects (Figure 3d, Figure S3c). SUV39H1 is potentially part of this subclass but was represented in the maintenance screen with only 2 sgRNAs and fell below the threshold of our filtering criteria. Polycomb proteins, chromatin remodeling factors, and RNA processing factors were also included in Class I, a subset of which were validated by generating individual lentiviral-mediated knockout mESC lines to assess their effects on maintenance (Figure 3d, Figure S3a and b). Class II genes were defined by comparing establishment and maintenance genes using a lower statistical cut-off for establishment (FDR<0.2, methods) (Figure 3e). These included several RNA processing factors such as Rbmxl1 (involved in heterochromatin compaction)^53^, and SFPQ (Splicing Factor Proline and Glutamine rich protein that has been shown to act as a co-repressor and to interact with NONO and RBM14^54, 55^, respectively) (Figure S3d). Other genes in this class include CDC45, a DNA replication factor and ARID4a, a chromatin remodeler (Figure S3d).

Notable factors within Class III included (1) the DNA/RNA helicase DHX9 and nine other RNA processing factors (Figure 3c, f, Figure S3c), (2) components of the DNA replication machinery WDHD1, RPA1, PRIM1, PRIM2, DNA2 and TIMELESS, and (3) several chromatin remodeling factors [including the nucleosome remodeling and deacetylase (NuRD) complex components, RBBP4 and CHD4, the BAF subunit PBRM1, HUSH and H3K9me3 interacting protein MORC2a, and the DNA replication independent histone chaperone HIRA] (Figure 3f, Figure S3c). In addition, our analysis revealed several other candidate genes that were less highly enriched class III members, including condensin components, co-repressor and DNA binding proteins, DNA damage and repair proteins, and nuclear periphery proteins Rae1 and NUP205 (Figure 3f, Figure S3e). These findings suggest that H3K9me3 heterochromatin maintenance in mammalian cells not only requires the core H3K9me and DNA methylation enzymes, but also depends on chromatin remodeling complexes, replisome-associated and nuclear periphery proteins, and RNA processing factors, including the conserved DHX9 helicase.

### Epigenetic maintenance of silencing is stabilized in differentiated cells

Although the TetR-KRAB-induced silent state could be maintained for multiple cell divisions in the absence of sequence-dependent initiation, it decayed over time in mESCs (Figure 1f). In fission yeast, DNA sequence-independent heterochromatin maintenance is opposed by the anti-silencing factor Epe1, which encodes a Jumonji (JmjC) family putative demethylase^7, 10^. Our genetic screen in mESCs showed that mutation of the JmjC domain-containing JMJD2c/KDM4c demethylase resulted in enhanced maintenance of the silent state but did not affect establishment (Figure 2b, Figure 3b, Figure 4a). In mESCs, H3K9 demethylases and 5mC hydroxylases are also highly expressed with roles in stem cell renewal, however their expression is reduced upon differentiation (Figure S4a)^56–59^. To test whether the maintenance of silencing was affected during the transition from pluripotency to a differentiated state, we first established the eGFP^-^ silent state in mESCs in -Dox medium and then differentiated them into neural progenitor cells (NPCs) in +Dox medium (Figure 4b, Figure S4b). Differentiation was validated by the reduction in expression of pluripotency genes and increase in expression of NPC marker genes using qRT-PCR (Figure S4b). Furthermore, we observed decreased expression of H3K9 demethylases, KDM4a and 4c, and 5mC hydroxylases, TET1 and TET2 in NPCs (Figure 4c). Throughout differentiation, cells were maintained in +Dox medium to ensure that the *8x-tetO* sites remained TetR-KRAB-free. Using FACS analysis, we quantified the fraction of the cells with silent or active eGFP reporter at different time points (Figure 4d). Strikingly, the silent state persisted in 100% of NPCs after 10 days (∼10 cell divisions) and 50 days (∼50 cell divisions). To test whether the differentiation process affected expression of the reporter gene, we differentiated mESCs into NPCs without first establishing the silent state and observed that almost all cells had an active reporter (Figure S4c). ChIP-qPCR experiments showed that an H3K9me3 domain established in mESCs was present after differentiation in NPCs cultured in -Dox medium (when TetR-KRAB is still recruited at the reporter locus) (Figure S4d), and was maintained in NPCs cultured in +Dox medium (that promotes the release of TetR-KRAB from the *tetO* sites) (Figure 4e). Additionally, bisulfite sequencing showed an increase in CG dinucleotide methylation in NPCs relative to mESCs with ∼96% of the CG dinucleotides methylated in NPCs and fully maintained following the release of TetR-KRAB (Figure 2f, Figure 4f). These results suggest that DNA sequence-independent epigenetic inheritance of heterochromatin is greatly stabilized in differentiated cells and that this stabilization correlates with increased levels of both H3K9me3 and DNA CpG methylation.

**Figure 4.**
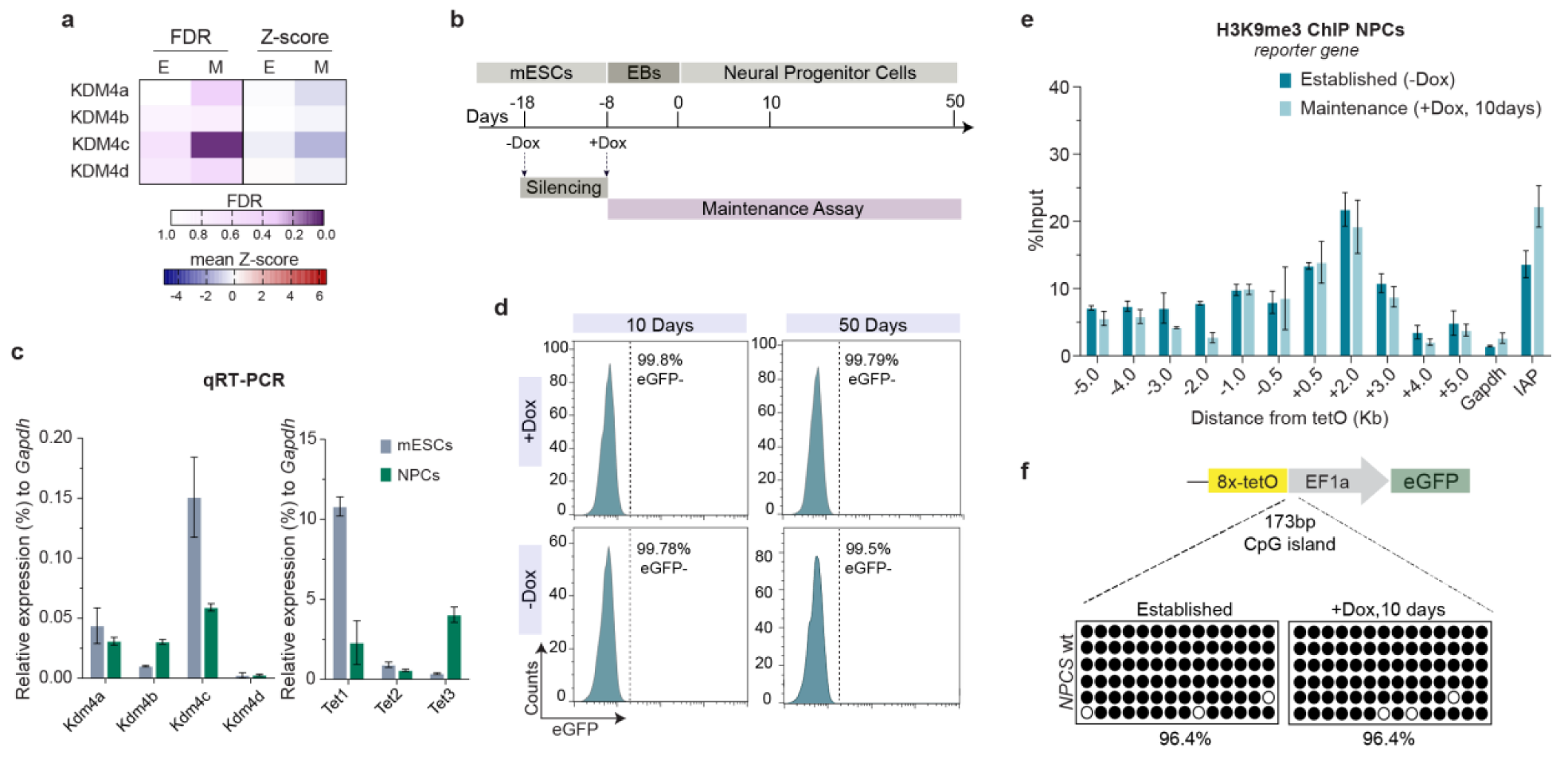
Epigenetic maintenance of silencing is stabilized upon differentiation. **a,** Heatmap depicting screen data for KDM4 demethylases. Gene-level FDR values and mean values of the Z-scores of all 5 sgRNAs targeting each gene in eGFP^+^ sorted versus unsorted cells during establishment (E) or maintenance (M) are shown. (n = two independent replicates). **b,** Schematic shows time-course of the differentiation of mESCs to embryonic bodies (EB) and finally to neural progenitor cells, and the outline of the assay used to test maintenance of silencing at the reporter locus in neural progenitor cells. **c,** qRT-PCR analysis showing expression levels of KDM4 demethylases and TET hydroxylases relative to *Gapdh* levels in mESCs and NPCs. Mean values are shown. Error bars, standard deviation (SD); n = 3 biological replicates. **d,** Flow cytometry histograms show eGFP expression in neural progenitor cells, that were differentiated from mESCs with a silenced reporter locus, at the indicated time points as shown in panel **c**, cultured either in medium with (+Dox) or without (-Dox) doxycycline. Percentages (%) indicate fraction of eGFP^-^ cells. **e,** ChIP-qPCR analysis for H3K9me3 at the *8x-tetO-eGFP* reporter locus and surrounding regions in neural progenitor cells (NPCs) cultured in the absence (Established, -Dox) or presence (Maintenance, +Dox) of doxycycline. *Gapdh* and *IAP*, are used as controls for euchromatin and heterochromatin H3K9me3 levels respectively. Values are shown as percentage (%) of input. Error bars, standard deviation (SD); n = 3 replicates. **f,** Bisulfite sequencing of DNA CpG methylation at the reporter locus in NPCs cultured in the absence (Established) or presence (+Dox) of doxycycline. Filled and open circles represent methylated and unmethylated CpGs, respectively. Percentages (%) indicate methylated CpGs.

### Deletion of *Dhx9* diminishes epigenetic maintenance

The DNA/RNA helicase DHX9 was among the top hits in our genetic screen required specifically for epigenetic inheritance of silencing. This and a previous report of DHX9 enrichment in heterochromatin fraction of mammalian cells prompted us to study its function in heterochromatin formation^53^. To investigate the role of DHX9 in epigenetic silencing, we mutated its gene with CRISPR-Cas9, resulting in a complete loss of protein (*Dhx9* KO) in mESCs, that had the inducible heterochromatin system and the eGFP reporter (Figure S5a). We then tested whether *Dhx9* deletion affected establishment and maintenance of silencing in the inducible reporter system. Consistent with its identification as a maintenance-specific factor in our screen, silencing was established successfully in *Dhx9* KO cells cultured in -Dox medium but with slower kinetics (Figure S5b). However, maintenance of the established silent state was defective (Figure 5a). While in wild-type cells, eGFP silencing persisted in ∼85% and ∼70% of cells 3 and 5 days after growth in +Dox medium, in *Dhx9* KO cells, eGFP silencing persisted in 53%-64% and 33%-44% at the corresponding time points, indicating more rapid decay (Figure 5a). *Dhx9* KO cells also had a slower duplication time of 20 hours compared to 14 hours for wild-type cells (Figure S5c). Therefore, the *Dhx9* KO maintenance defect is likely to be more robust when comparing corresponding cell division numbers rather than at absolute time points. For comparison, we generated clonal lines with disruption of *Uhrf1*, a top class I hit in our screen, which encodes an E3 ubiquitin ligase that interacts with the CpG methyltransferase, DNMT1, and is required for DNA methylation maintenance (Figure S5d)^60, 61^. Although different clones of *Uhrf1* KO cells showed a moderate to weak effect in the establishment of silencing at the reporter locus, only 9-11% maintained silencing 3 days after growth in +Dox medium, a dramatic 8-fold reduction relative to wild-type cells (Figure S5e). Using ChIP-seq and ChIP-qPCR, we compared the effects of *Dhx9* or *Uhrf1* deletion on H3K9me3 establishment and maintenance at the reporter locus. We found that during establishment, the reporter locus showed diminished enrichment of H3K9me3 in *Dhx9* KO cells but not in *Uhrf1* KO cells (Figure 5b, c, Figure S5f). Correspondingly, CpG methylation at the reporter locus was reduced in both *Dhx9* KO (46%) and *Uhrf1* KO (23%) cells compared to wild-type cells (74%) (Figure 5d). In the maintenance phase, H3K9me3 was further depleted in *Dhx9* KO cells and lost in *Uhrf1* KO cells indicating that DNA CpG methylation was required for epigenetic inheritance of H3K9me3 (Figure 5c, Figure S5g). Together, these results uncovered a novel role of DHX9 in H3K9me inheritance and the propagation of silencing and highlight the importance of DNA methylation in maintaining H3K9me-dependent silencing.

**Figure 5.**
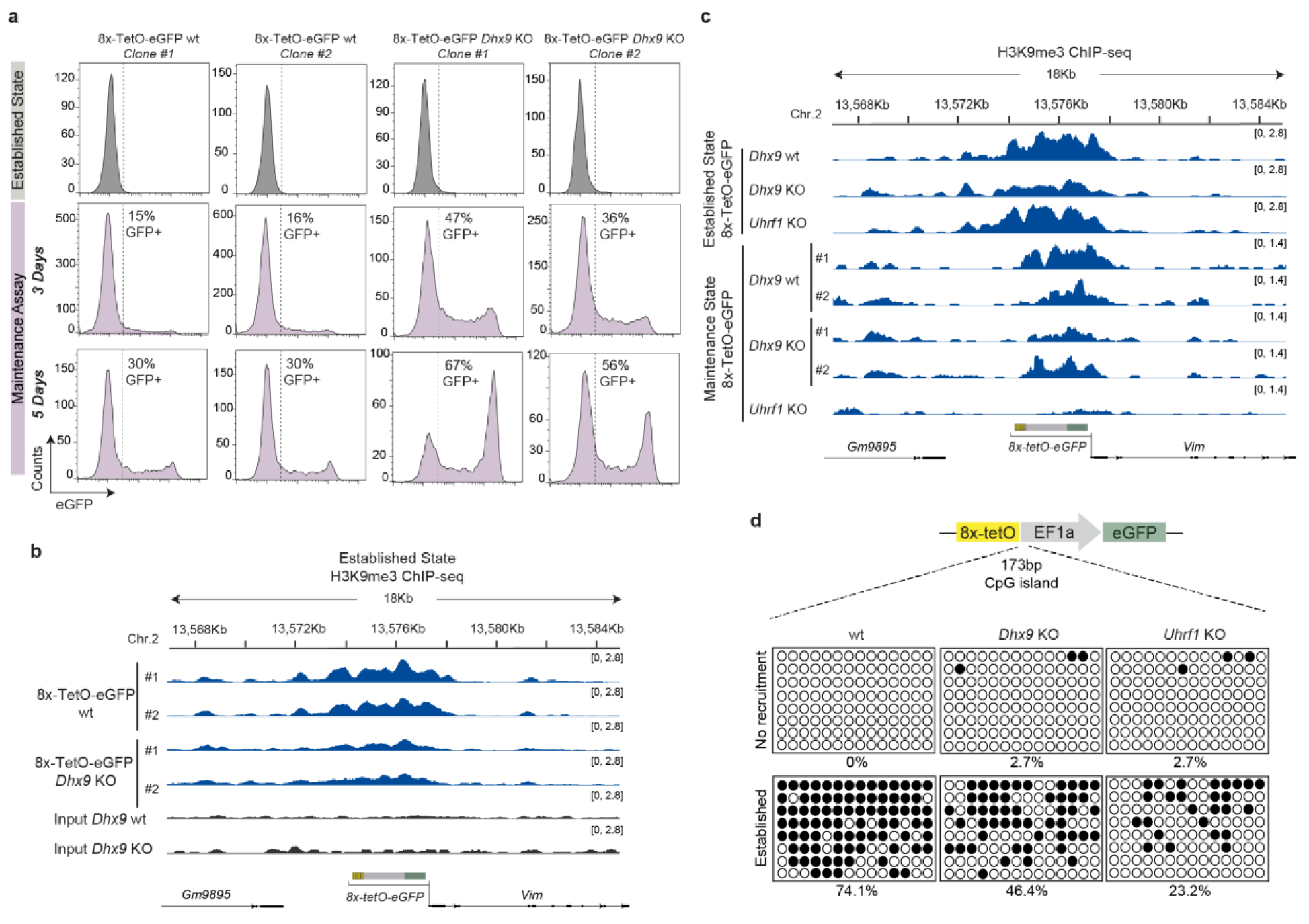
*Dhx9* deletion results in defective maintenance of silencing. **a,** Flow cytometry histograms showing eGFP expression in *8x-tetO-eGFP* wild-type (wt) and *Dhx9* KO mESCs cultured in medium without doxycycline for 12 days (Established State, grey) and three (DIV3, pink) and five (DIV5, blue) days after adding back doxycycline to assess maintenance. Percentages (%) indicate fraction of eGFP^+^ cells. **b,** Genome tracks of input and H3K9me3 ChIP-seq for regions surrounding the reporter locus after establishment of silencing in *8x-tetO-eGFP* wild-type (wt) and *Dhx9* KO mESCs. Normalized reads are presented in brackets. Top, chromosome coordinates. **c,** Genome tracks of H3K9me3 ChIP-seq for regions surrounding the reporter locus right before (Established State) and after adding back doxycycline for five days to assess the maintenance in *8x-tetO-eGFP* wild-type (wt), *Dhx9* KO, and *Uhrf1* KO mESCs. Normalized reads are presented in brackets. Top, chromosome coordinates. **d,** Bisulfite sequencing of DNA CpG methylation at the reporter locus in *8x-tetO-eGFP* wild-type (wt), *Dhx9* KO, and *Uhrf1* KO mESCs cultured before establishment in the presence (No recruitment) or after establishment in the absence (Established) of doxycycline. Filled and open circles represent methylated and unmethylated CpGs, respectively. Percentages (%) indicate methylated CpGs.

### Dhx9 is required for H3K9 methylation and silencing at Major Satellite Repeats

To investigate the role of DHX9 in H3K9me3 maintenance at endogenous heterochromatic regions, we performed H3K9me3 ChIP-seq in wild-type and *Dhx9* KO mESCs. Genomic regions with H3K9me3 were annotated using Epic2 peak caller^62^. Both wild-type and *Dhx9* KO samples had a median H3K9me3 peak width of ∼2500 bp (Figure 6a). To identify differentially methylated peaks in *Dhx9* KO, we used a statistical package, DiffBind,^63^ which identified 89,055 consensus peaks for quantitative analysis (Supplementary Table 5). Most of these peaks were localized at intergenic (63% of the 89,055 peaks) or intronic regions (32%), and >95% of the peaks overlapped with annotated repeat elements (Figure 6b, c). Broadly, *Dhx9* KO showed reduced H3K9me3 enrichment in peaks individually (Fig 6c) and collectively (Figure S6a). The vast majority of significantly altered H3K9me3 peaks (n=648) showed decreased H3K9me3 enrichment (90%, 585/648) with only a small fraction showing increased (10%, 63/648) (Figure 6c). Peaks with increased H3K9me3 in *Dhx9* KO corresponded mainly to regions with relatively low H3K9me3 signal (Figure 6c).

**Figure 6.**
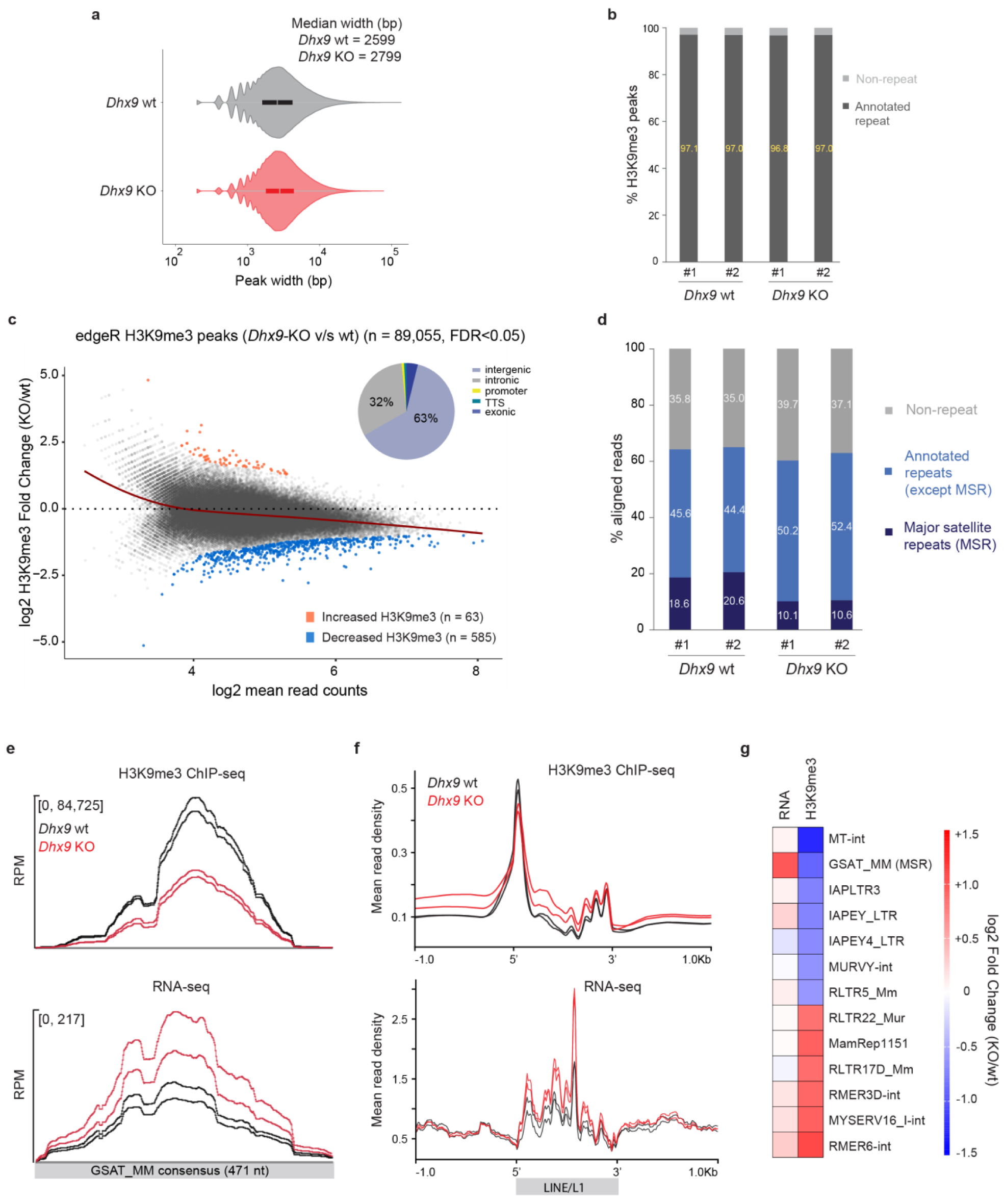
DHX9 is required for H3K9 methylation and silencing at the major satellite repeats. **a,** Violin plots showing the distribution of H3K9me3 peak width in *8x-tetO-eGFP* wild-type (wt) and *Dhx9* KO mESCs. **b,** Plots displaying the percentage of H3K9me3 peaks overlapping with annotated repeats in wild-type and *Dhx9* KO mESCs. **c,** Scatterplot showing the mean H3K9me3 read counts (x-axis) in all samples for the 89,055 consensus peaks identified by DiffBind and log_2_ ratio of H3K9me3 enrichment in *Dhx9* KO (KO) versus wild-type (wt) mESCs (y-axis). H3K9me3 peaks with increased (orange) and decreased (blue) signal in *Dhx9* KO are shown at an FDR < 0.05. A LOESS local regression fit (red line) across the scatterplot shows a trend for overall decreased methylation in *Dhx9* KO cells. Source data for this figure are provided in Supplementary Table 5. Inset: Pie chart showing genomic annotation of H3K9me3 peaks. **d,** Plots displaying the percentage of H3K9me3 reads overlapping with annotated repeats, major satellite repeats (MSRs, GSAT_MM), and non-repeat regions in wild-type and *Dhx9* KO mESCs. **e,** Plots showing the normalized average density of H3K9me3 ChIP-seq (top) and RNA-seq reads (bottom) across MSR consensus sequence in wild-type (wt) and *Dhx9* KO mESCs. **f,** Same as in **e** for LINE/L1 repeat family. **g,** Heatmap displaying the fold change of the RNA levels (left) and H3K9me3 enrichment (right) of the top H3K9me3 decreasing and top H3K9me3 increasing repeat families in *Dhx9* KO (KO) versus wild-type (wt) mESCs. Source data for this figure are provided in Supplementary Table 6.

Since most H3K9me3 peaks overlap with repetitive genomic regions, we asked how *Dhx9* deletion affected H3K9me3 and transcriptional silencing of annotated repeat elements. By comparing the total number of H3K9me3 ChIP-seq reads aligning to each annotated repeat element in *Dhx9* KO and wild-type cells (Supplementary Table 6), we found that Major Satellite Repeats (MSRs, annotated as GSAT_MM) were highly enriched for H3K9me3 in wild-type cells (∼20% of the reads align to GSAT_MM; Figure 6d) but dramatically decreased in *Dhx9* KO cells (Figure 6d, e, Figure S6b). H3K9me3 ChIP-qPCR further confirmed this observation (Figure S6d). In a similar context, the characteristic H3K9me3 enrichment on the 5′ ends of intact LINE/L1 repeats^64, 65^ was reduced in *Dhx9* KO cells, but increased on LINE/L1 body and flanking regions (Figure 6f). An elevated H3K9me3 signal was also observed more generally at regions distal from peak centers when aggregating the signal across all peaks (Figure S6a), consistent with the idea that the loss of H3K9me3 at the peaks may lead to a genome-wide increase in H3K9me3. Other repeat families that showed a pronounced reduction in H3K9me3 in *Dhx9* KO relative to wild-type mESCs included endogenous retroviral repeats (ERVs), such as ERVL-MaLR repeat (MT-int), ERV1 (MURVY-int, RLTR5_Mm), ERVK (IAPLTR3, IAPEY_LTR, IAPEY4_LTR) repeats, and the Pol I-transcribed region (LSU-rRNA, SSU-rRNA) but not intergenic spacer region of rDNA repeats (Figure S6b, data not shown).

Loss of H3K9me3 at MSRs and LINE/L1 repeats in *Dhx9* KO corresponded with increased RNA levels, measured by rRNA-depleted total RNA-seq (Figure 6e, f, Figure S6c), suggesting that DHX9 is required for H3K9me3 maintenance and silencing at these repeat regions. However, reduction in H3K9me3 after DHX9 depletion was not always accompanied by a change in RNA abundance (e.g., MT-int, IAPLTR3, MURVY-int, RLTR5_Mm), suggesting that the decrease in H3K9me3 levels was insufficient to derepress these repeats or transcriptional activation mechanisms were absent in mESCs (Figure 6g, Figure S6c). Some ERVK repeats showed an increase in H3K9me3 signal but no change in RNA levels in *Dhx9* KO cells (Figure 6g, Figure S6c, Supplementary Table 6). This could be attributed to the function of KRAB-ZNFs, which were shown to repress ERVs^66^ and were upregulated in *Dhx9* KO cells (Figure S7a). Overall, our findings uncover a role of DHX9 in the regulation of H3K9me3 and transcriptional silencing, predominantly at MSRs.

Although most of the peaks that lose H3K9me3 in *Dhx9* KO overlapped with repeat elements, a small subset (6%, 38/585) overlapped gene promoters (TSS +/-2kb, Supplementary Table 5). Most changes in H3K9me3 at promoters did not affect genic RNA levels with 4 exceptions (Figure S6b). Loss of H3K9me3 at *Nanog* and *Trim52* promoters corresponded with increased RNA abundance, indicating transcriptional de-repression (Figure S6b, c). By contrast, H3K9me3 loss at *Ngln2* and *Pcdhgb4* showed decreased RNA levels (Figure S6b, d). In addition to H3K9me3 promoter peaks, these two genes contained H3K9me3 within their gene bodies. Reduction of gene body H3K9me3 could result in defective silencing of cryptic promoters that could interfere with proper transcription of the gene (Figure S6d). Although many genes showed significant changes in RNA levels in *Dhx9* KO mESCs (Supplementary Table 7), only a few had H3K9me3 at their promoters (4%, 15/375 of upregulated genes; 13%, 31/242 of downregulated genes) (Figure S6a, Supplementary Table 6). Among the genes with increased RNA levels in *Dhx9* KO mESCs, KRAB-containing zing-finger (KRAB-ZNFs) genes were enriched (n=54/375, Figure S6a). H3K9me3 around the KRAB-ZNFs gene clusters remained unaltered in *Dhx9* KO cells (Figure S6e), suggesting an H3K9me3-independent role of DHX9 in KRAB-ZNF gene repression. Overall, these findings show that DHX9 is required for maintenance of H3K9me3 and silencing at MSRs and other repeats and some euchromatic genes and contributes to repression of ZNF family of transcription factors by a mechanism that may be independent of H3K9me3.

### DHX9 loss results in accumulation of heterochromatin-associated RNAs and increase of RNA/DNA hybrids at MSR

Previous studies have shown that the DHX9 helicase interacts with RNA/DNA hybrids and prevents R-loop accumulation in mammalian cells^33, 67^. R-loops are structures created in genomic regions where duplexes of RNA with its complementary single-stranded DNA are formed while the second DNA strand is displaced^68^. R-loops have the potential to form over a large proportion of the genome, including pericentric heterochromatin (localized at major satellite repeats), as well as in some interspersed repeats in the genome^69, 70^. The DHX9 helicase resolves R-loops likely by unwinding RNA/DNA hybrids. Thus, we hypothesized that the loss of heterochromatin maintenance in *Dhx9* KO cells might result from the accumulation of site-specific R-loops. We first asked whether DHX9 was associated with chromatin to test this hypothesis. Biochemical isolation of the nucleoplasmic and chromatin fractions in mESCs showed that DHX9 was present in both fractions (Figure S8a). To test whether DHX9 was recruited to the reporter locus in a heterochromatin-dependent manner, we conducted ChIP-qPCR. We found that DHX9 recruitment was increased after the establishment of silencing (Figure 7a), suggesting that heterochromatin formation promotes DHX9 recruitment. Moreover, ChIP-qPCR showed DHX9 was enriched at MSRs, suggesting that it may have a direct role in heterochromatin regulation at endogenous repeats (Figure 7b).

**Figure 7.**
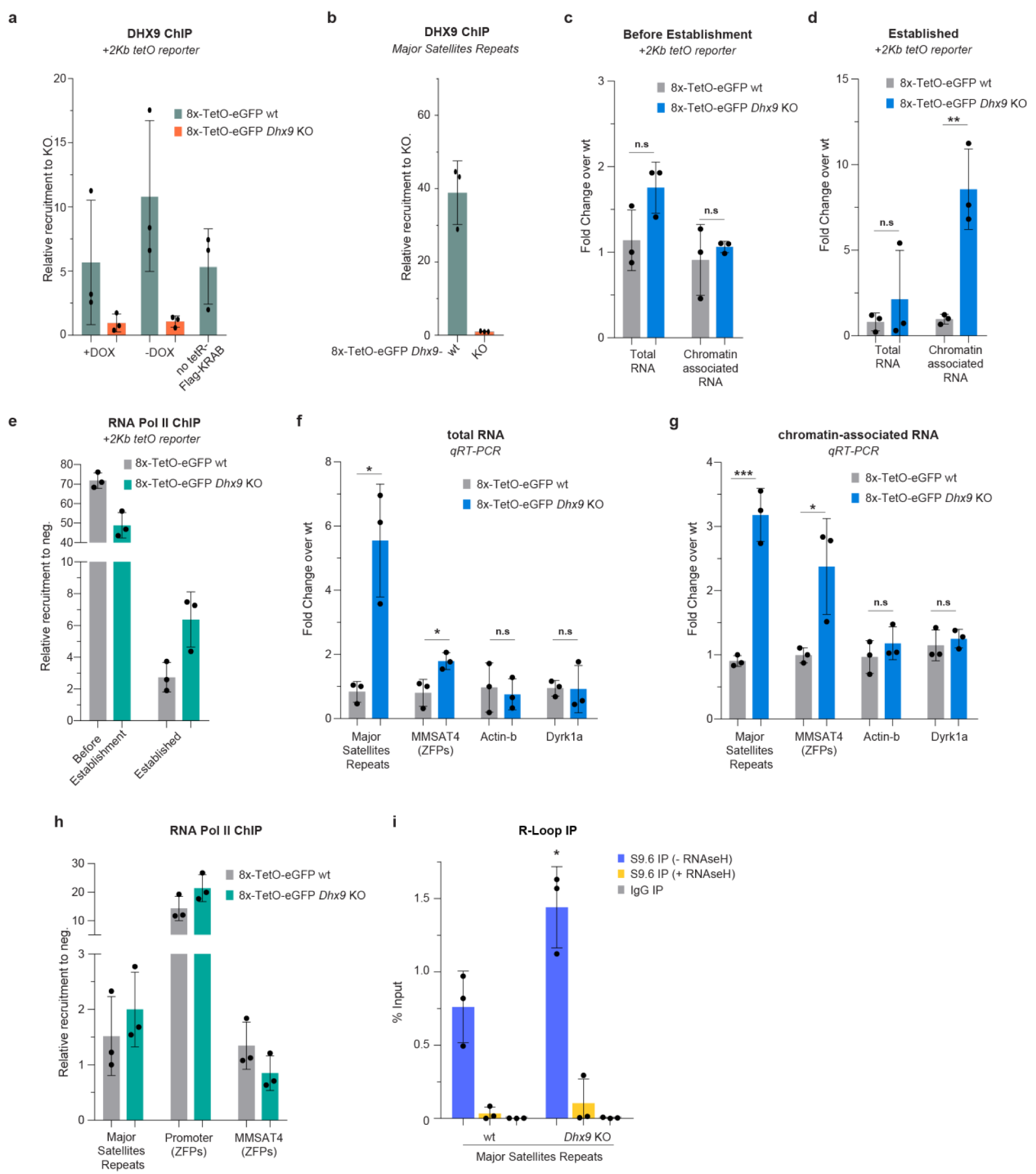
DHX9 loss results in accumulation of heterochromatin-associated RNAs and increased RNA/DNA hybrids at MSR. **a,** ChIP-qPCR analysis for DHX9 enrichment at the *8x-tetO-eGFP* reporter locus in wild-type (wt) and *Dhx9* KO mESCs before establishment (+Dox) or after establishment of silencing (-Dox), and in *8x-tetO-eGFP* mESCs without TetR-Flag-KRAB. Values are shown as relative recruitment over KO cells. Error bars, standard deviation (SD); n = 3 biological replicates. **b,** ChIP-qPCR analysis for DHX9 enrichment at major satellite repeats (MSRs) in wild-type (wt), *Dhx9* KO, and in *8x-tetO-eGFP* mESCs without TetR-Flag-KRAB. Values are shown as relative recruitment over KO cells. Error bars, standard deviation (SD); n = 3 biological replicates. **c,** qRT-PCR analysis showing total and chromatin-associated RNAs of the *8x-tetO-eGFP* reporter gene in wild-type (wt) and *Dhx9* KO mESCs before establishment. Values are shown as fold change over *wt* cells. Error bars, standard deviation (SD); n = 3 biological replicates. **d,** Same as in (**c**) after establishment of silencing. **e,** ChIP-qPCR analysis for RNA Pol II binding at the *8x-tetO-eGFP* reporter locus in wild-type (wt) and *Dhx9* KO mESCs before establishment and after establishment of silencing. Values are shown as relative recruitment over a negative region in chromosome 1. Error bars, standard deviation (SD); n = 3 biological replicates. **f,** Same as in **c** showing total RNA changes at major satellite repeats and MMSAT4 elements in *8x-tetO-eGFP* wild-type (wt) and *Dhx9* KO mESCs. **g,** Same as in **f** showing chromatin-associated RNAs. **h,** Same as in **e** displaying RNA Pol II binding at the indicated regions. **i,** DNA-RNA immunoprecipitation (DRIP) analysis using the RNA-DNA hybrid antibody S9.6 at the major satellite repeats in wild-type (wt) and *Dhx9* KO mESCs. Treatment with RNAse H, and immunoprecipitation with IgG control antibody was used as negative controls for specificity. Error bars, standard deviation (SD); n = 3 biological replicates. *P*-values in all panels calculated with pairwise unpaired t tests are indicated with asterisks. n.s., not significant (*P* > 0.5), *, *P* < 0.05, **, *P* < 0.01, ***, *P* < 0.001.

To investigate whether DHX9 effects on heterochromatin were mediated via the regulation of heterochromatin-bound RNAs, we isolated the chromatin fraction in *Dhx9* KO mESCs and quantified the levels of the associated RNAs by qRT-PCR. First, we tested the effects of *Dhx9* KO on the levels of steady-state total and chromatin-associated RNAs transcribed from the *8x-tetO* reporter locus. Before the establishment of silencing, we did not detect any change in *egfp* reporter RNA levels in the *Dhx9* KO cells relative to wild-type cells (Figure 7c). Following establishment, although we did not observe an alteration in the steady-state total RNA levels, we observed a marked increase in the chromatin-associated *egfp* RNA levels in *Dhx9* KO cells (Figure 7d). As expected, RNA polymerase II (Pol II) ChIP-qPCR showed that RNA Pol II occupancy was reduced following the establishment of silencing, but to a lesser extent in *Dhx9* KO relative to wild-type cells (Figure 7e). Together, these findings suggest that DHX9 promotes the processing or release of repeat-associated heterochromatic RNAs.

We next used qRT-PCR to verify the RNA-seq data on the de-repression of MSRs and the MMSAT4/ZNF repeats. We detected a significant increase in total RNA levels of MSR and MMSAT4/ZNF regions in *Dhx9* KO cells (Figure 7f). As with the reporter locus (Figure 7d), heterochromatin-associated RNAs at these regions were increased in *Dhx9* KO cells, suggesting that DHX9 loss resulted in their accumulation at the transcriptional or co-transcriptional level (Figure 7g). The increased accumulation of MSR and MMSAT4 chromatin-bound RNAs was not accompanied by increased RNA Pol II occupancy at the MSR repeats, the MMSAT4 repeats, or the promoter of ZNFs genes (Figure 7h), suggesting that *Dhx9* KO did not affect the levels of transcription from the repeats. Since DHX9 has been implicated in unwinding RNA/DNA hybrids and inhibits R-loop accumulation, the increase in chromatin-associated RNAs in *Dhx9* KO cells may result from defective R-loop processing at heterochromatic loci. We tested this hypothesis by performing DNA-RNA immunoprecipitation (DRIP) analysis in *Dhx9* KO versus wild-type mESCs. Native nuclear extracts were immunoprecipitated (IP) with the RNA-DNA-hybrid-specific antibody, S9.6, and the purified DNA was used for qPCR analysis. RNAse H1 treatment of parallel IPs was used to control for R-loop-specific precipitation. It was technically challenging to detect R-loops at the reporter locus since the establishment of silencing leads to low levels of RNA and a poor signal-to-noise ratio. However, we observed increased R-loop levels at the endogenous heterochromatic major satellite repeats in *Dhx9* KO cells (Figure 7i). Together, these results suggest that DHX9 is required for resolving RNA/DNA hybrids at heterochromatic loci, such as MSRs, and that the accumulation of RNA/DNA hybrids leads to defects in heterochromatin maintenance.

## Discussion

Our findings suggest that efficient heterochromatin establishment requires the combined action of multiple H3K9 methyltransferases and downstream effector proteins and uncover an unexpected role for DNA methylation in both the establishment and maintenance of H3K9me-dependent silencing (Figure 8a, b). We have demonstrated that inheritance of H3K9me-dependent heterochromatin is independent of the underlying DNA sequence, and although it is metastable in mESCs, it becomes remarkably stable upon their differentiation to NPCs. In mammalian cells, the coupling of read and write positive feedback loops associated with each H3K9me and 5mC may allow DNA sequence-independent epigenetic inheritance of heterochromatin, which in organisms lacking DNA methylation may usually require input from specific DNA sequences or RNAi. The reciprocal reinforcement of the two feedback loops likely increases the rate of re-establishment of silent domains during DNA replication and counteracts activities that remove histone modifications. In agreement with this hypothesis, we found that increased stability of heterochromatin inheritance in differentiated cells, which correlates with increased levels of H3K9me and 5mC, results from the downregulation of enzymes that erase H3K9me and 5mC. The shift in the balance between H3K9me and 5mC deposition and removal during development may therefore act as a key event that determines the stability of mammalian heterochromatin^59^ and suggests deep conservation of heterochromatin maintenance pathways from fission yeast to mammals^7^. In addition, our findings uncover a role for the DHX9 R-loop resolvase in safeguarding heterochromatin stability against chromatin remodeling events that are associated with replication stress and DNA damage repair.

**Figure 8.**
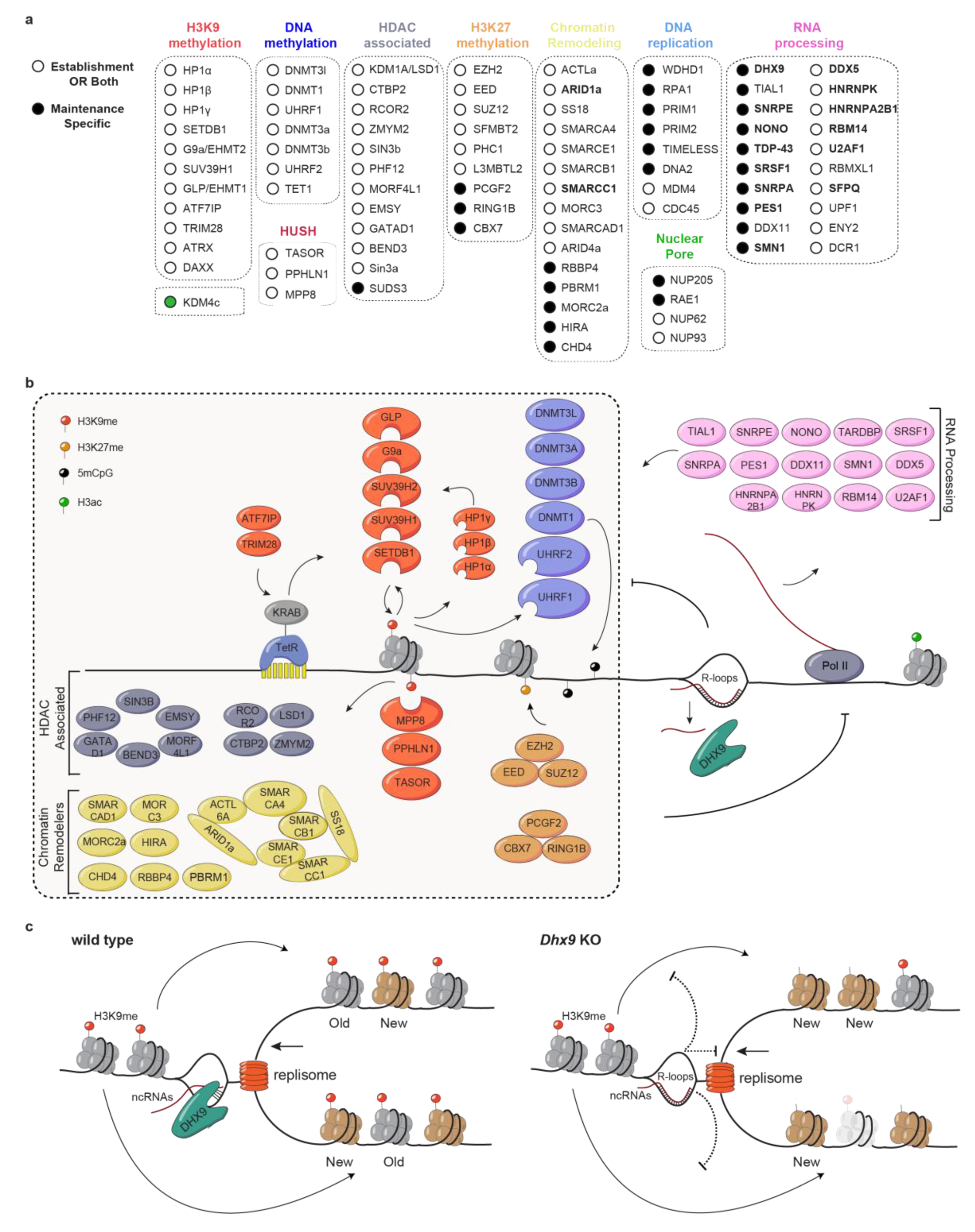
Requirements for the establishment and epigenetic inheritance of H3K9 methylation. **a,** Summary of representative proteins required for establishment and maintenance (white circles) and proteins required solely for maintenance (black circles) of H3K9me domains. Note that KDM4c (green circle) mutation enhances maintenance. Factors implicated in R-loop metabolism, including those with defects in splicing, are highlighted in bold. **b,** Schematic diagram highlighting the molecular activities required for H3K9me-dependent silencing, depicting representative genes. **c,** Diagram illustrating the proposed role of the DHX9 helicase in heterochromatin maintenance. Increased R-loop formation (associated with RNA polymerase II transcription, not shown) in *Dhx9* KO cells inhibits replisome progression leading to replication stress and local loss of H3K9me nucleosomes.

Our analysis revealed that, in addition to SETDB1, the other H3K9 methyltransferases G9a, GLP, SUV39H1, and SUV39H2, but not SETDB2, are required for the establishment of *de novo* heterochromatin at the ectopic locus. Previous studies have shown that multiple H3K9 methyltransferases (SUV39H1, G9a, GLP, SETDB1) can associate together in a multimeric protein complex, however, whether they functionally cooperate for silencing of a single locus had not been previously investigated^71^. Although the molecular basis for the requirement of multiple H3K9 methyltransferases (this study) remains unknown, likely possibilities include their roles in the transition from H3K9me1 to H3K9me3 or their distinct read-write abilities that may be required for H3K9me spreading. In good agreement with previous reports, we also observe TetR-KRAB-induced DNA methylation at the reporter locus. However, the requirement of multiple DNA methyltransferases and their interacting partners for the establishment of H3K9me-dependent silencing was unexpected^72^ (Figure 8a, b). We provide evidence that reduced silencing resulting from the loss of DNMTs or UHRF1 stems from reduced DNA methylation. Since the loss of DNMTs or UHRF1 does not affect H3K9me3 during establishment of silencing but is accompanied by rapid decay of H3K9me3 during maintenance; we conclude that DNA methylation plays critical roles in both phases of silencing. Furthermore, this observation suggests that in mammalian cells, H3K9me3 by itself is not heritable, despite numerous read-write modules present in the methyltransferases themselves or imparted through their association with HP1 proteins. UHRF1 binds to methylated CpG dinucleotides and H3K9me3 through distinct domains, providing a physical and functional link between H3K9me and DNA methylation, and promotes the ubiquitination of H3 on lysines 18 and 23 to promote DNMT1 recruitment for maintenance of DNA methylation^60, 61, 73–78^. The requirement for UHRF1 in maintenance of both DNA methylation and H3K9me may be explained by its reported ability to associate with G9a and SUV39H1^79^, creating reciprocal reinforcing positive feedback loops between DNA methylation and H3K9me3 – an emerging principle in heritable epigenetic modifications^80^.

In addition to components of H3K9me and DNA methylation pathways, we show that heterochromatin establishment and maintenance require the HUSH complex, HDAC-associated proteins, chromatin remodeling proteins (mainly components of the BAF complex), nuclear pore complex (NPC) subunits and Polycomb group proteins (Figure 8a, b). Despite evidence on the role of these proteins in heterochromatin formation and silencing, the molecular mechanisms underlying their functions in heterochromatin establishment and maintenance are not understood^23, 45–48, 65, 81–86^. Accumulating evidence highlights the function of the HUSH complex in initiating H3K9me and silencing long interspersed element-1 retrotransposons (L1s), endogenous retroviruses and long intronless transgene RNAs^65, 81, 82, 87^. Our findings indicate that the HUSH complex also acts downstream of H3K9me since H3K9me at our reporter locus is initiated by TetR-KRAB binding. Moreover, previous studies in *S. pombe* showed that Rae1 and NUP205 (NPC subunits) are enriched in Swi6/HP1 heterochromatin fraction, and NUP93 is required for clustering of heterochromatic foci and epigenetic inheritance of heterochromatin^88^(Figure 8a). Our data suggest that their role(s) in heterochromatin formation and silencing may be conserved in mammalian cells. Finally, the strong requirement for Polycomb proteins in establishment and maintenance of an H3K9me-dependent silencing is notable. Although co-enrichment of H3K9me and H3K27me has been previously reported on some LINE elements, and it has also been proposed that DNMTs recognize and target PRC-modified regions^41–44^, the functional importance of these interactions in silencing has remained unclear. Our findings raise the possibility that PRC1/2 complexes contribute to maturation of H3K9me heterochromatic domains, for example, by binding to CpG islands prior to their methylation by DNMTs.

Our screen and analyses also uncover novel factors and molecular pathways required solely for heterochromatin maintenance, including several DNA replication and RNA processing factors (Figure 8a, b). Our identification of replisome proteins as maintenance-specific factors is consistent with the central role of the replication machinery in parental histone transfer and epigenetic inheritance^89^. For example, replication protein A (RPA) (Figure 8a) binds single-stranded DNA during DNA replication and has a role in replication-coupled nucleosome assembly^90^. RPA has been reported to bind the histone chaperone HIRA^91^, another maintenance-specific factor found in our screen, suggesting that it may also play a role in DNA replication-independent nucleosome assembly. Another replication protein, WD Repeat and HMG-Box DNA Binding Protein 1 (WDHD1), identified with a role in heterochromatin maintenance in this study, has been implicated in the formation of pericentric heterochromatin in mammalian cells^92^. Importantly, a previous study identified mutations in replisome proteins that result in defective heterochromatin maintenance in fission yeast *S. pombe*^29^, several of which were also revealed by our screen. Together with previous studies, our results suggest that the DNA replication machinery plays a conserved role in epigenetic inheritance of heterochromatin.

Among the RNA processing proteins identified in this study, we found that deletion of DHX9, an RNA/DNA helicase, results in defective heterochromatin maintenance and is associated with increased transcription at major satellite repeats and accumulation of R-loops. DHX9 is a member of the DExH-box family and SF2 superfamily of helicases that binds to single-stranded DNA and RNA but preferentially unwinds RNA/DNA hybrids in RNA displacement loops (R-loops) and has also been shown that resolves RNA secondary structures^67, 93–95^. Consistent with these observations, DHX9 is strongly enriched in the RNA/DNA hybrid interactome in human cells and promotes R-loop suppression and transcription termination^33^. RNA/DNA hybrids can form at several genomic regions, including the mammalian pericentric heterochromatin (MSRs). Indeed, major satellite repeat transcripts are chromatin-associated and have a secondary structure that facilitates RNA/DNA hybrid formation^25^. Accumulation of R-loops is associated with stalling of replication forks and leads to replication stress^96–98^. DHX9 has been reported to resolve R-loops to prevent replication stress and its loss leads to increased 53BP1 foci and other events associated with DNA repair^33, 34^. In particular, DHX9 is required for replication restart from stalled replication forks^99^, which may prevent nucleosome loss and remodeling associated with DNA damage repair. We therefore propose that the accumulation of heterochromatin-associated R-loops in *Dhx9* KO cells leads to replication stress, which may then lead to local loss of H3K9me3 nucleosomes (Figure 8c). This loss of heterochromatin appears to occur in a time-dependent manner during heterochromatin maintenance, suggesting that in the absence of DHX9, heterochromatin may only be compromised following stochastic rounds of transcription and R-loop formation. Our findings suggest that in addition to its well-established role in promoting genome instability, replication stress can lead to epigenetic instability. In addition, our results raise the possibility that some of the disease-associated phenotypes in Dhx9 mutants may result from heterochromatin loss^100^.

## Materials and Methods

### Cell Culture

E14TG2a mES cells (ATCC, #CRL-1821) were cultured on 0.1% gelatin-coated (Sigma, #G1890) plates in Dulbecco’s modified Eagle’s medium (DMEM) supplemented with 15% heat-inactivated fetal bovine serum (FBS) (Gibco, #10437), 1% MEM non-essential amino acid (Gibco, #11140), 2 mM Glutamine (Gibco, #25030081), 100 μM β-mercaptoethanol (Sigma, #63689), 10 U/ml penicillin/streptomycin (Gibco, #15140), 1,000 U/ml LIF (Stemcell, #78056) at 37 °C with 5% CO2. 293FT cells (Gibco, #R70007) used for the production of lentiviruses were maintained in DMEM with 10% heat-inactivated FBS, 2 mM glutamine and 100 U/ml penicillin/streptomycin following standard culture conditions. Differentiation of mESCs to neural progenitor cells (NPCs) was performed as previously described with some modifications^101^. Four days after the onset of embryonic body (EB) formation, we added 2 μM retinoic acid (Sigma R-2625). After dissociation of EBs, cells were seeded on 0.1% gelatin-coated plates and NPCs are cultured in N2B27 medium (1:1, Neurobasal:DMEM-F12, supplemented with N2 and B27, ThermoFisher) in the presence of 10 ng/ml bFGF (Peprotech) and 10 ng/ml EGF (Peprotech) ^102^. NPCs were dissociated gently with accutase (Invitrogen) for passaging and cultured in 5 μg/ml doxycycline when indicated.

### Generation of the reporter mESCs line

The *8x-tetO-eGFP* reporter construct was generated as follows: ∼800bp homology arms (flanking both sides at the integration site, found at position +37bp in NM011701 (*Vimentin* gene)) were PCR amplified from gblocks gene fragments synthesized at Integrated DNA Technologies (IDT). PCR amplified *8xtetO-EF1a* promoter-eGFP cassette from iDuet101a (Addgene, #17629) was cloned with Gibson assembly in frame with the SV40pA sequences that were PCR amplified from lentiCRIPSRv2 (Addgene, #52961). Gibson cloning was subsequently used to simultaneously encompass digested *8xtetO-EF1a* promoter-eGFP-SV40pA cassette in frame with the homology arms and the whole insert was cloned into the pSMART-HCKAN (Lucigen) vector, assembling the donor plasmid. To integrate the reporter gene in the genome, homology-dependent recombination mediated by CRISPR-Cas9 genome editing was induced via a spCas9/sgRNA ribonucleoprotein (RNP) complex reconstituted *in vitro* before insertion into the cells. sgRNA (5′-TGTGAATCGTAGGAGCGCTG-3′) was synthesized via *in vitro* GeneArt™ Precision gRNA Synthesis Kit (ThermoFisher, A29377) following manufacturer’s instructions. SpCas9 protein was purified by the Initiative for Genome Editing and Neurodegeneration Core in the Department of Cell Biology at Harvard Medical School. Donor plasmid, and spCas9/sgRNA RNP were delivered to cells by electroporation with Neon transfection system (ThermoFisher, MPK1025). Single eGFP^+^ cells were FACS sorted and seeded in 96-well plates. Single clones were picked, expanded, and screened for insertion by PCR. Validation of positive clones was performed by sanger sequencing.

*8xtetO-EF1a-eGFP* reporter mESC line was further engineered to harbor TetR-3xFlag-KRAB protein or TetR-3xFlag inserted at the *Rosa26* locus. The constructs for these lines were generated as follows: ∼650-750bp homology arms (flanking both sides at the integration site, found at position chr6:113,076,053 (GRCm38/mm10) (Gt(ROSA)26Sor) were PCR amplified from gblocks gene fragments synthesized at IDT. First, we generated PCR amplified EF1a promoter and TetR-NLS signal cassette from pLV-tTRKRAB-red (Addgene, #12250). 3xFlag-KRAB was PCR amplified from the same vector with a forward primer including the sequence encoding 3xFlag and SV40pA amplified from lentiCRIPSRv2. The inserts were cloned with Gibson assembly in frame with a neomycin-HSV tk pA cassette and the homology arms (HomL or HomR). The whole cassette (HomL-EF1a-TetR-3xFlag-<u>KRAB</u>-SV40pA-NeoR-HomR) with or without KRAB was cloned into the pSMART-HCKAN (Lucigen) vector, assembling the donor plasmid. To integrate the above cassettes in the genome the same method was used as above using sgRNA sequence (5′-ACTGGAGTTGCAGATCACGA-3′) that targets intron1 of the *Rosa26* locus. Single cells were FACS sorted and selected for neomycin resistance. Single clones were picked, expanded, and screened for insertion by PCR. Validation of positive clones was performed by sanger sequencing and western blot.

### EpiChromo Library generation

Single guide RNAs (sgRNAs) targeting 1160 genes including all known chromatin and epigenetic regulators, DNA replication factors, nuclear periphery factors, RNA processing factors, and proteins identified in proteomic analyses of heterochromatin composition were designed to generate a pooled library, named EpiChromo (Supplementary Table 1). The optimized sgRNA design strategy and rule set scoring (Azimuth 2.0) to determine editing activity scores described earlier ^103^ (http://www.broadinstitute.org/rnai/public/analysis-tools/sgrna-design) were used as the basis for the design of EpiChromo sgRNAs. Top-5 ranked sgRNAs targeting the coding domains of each gene after on-target and off-target analysis were picked for library construction. We excluded sgRNAs with a BsmBI recognition site in their sequence to allow subcloning into the lentiCRISPRv2 plasmid after BsmB1 digestion. We included 150 control sgRNAs that do not target the mouse genome in the EpiChromo library resulting in a total of 5,950 sgRNAs (Supplementary Table 2). EpiChromo library is also available at the Gene Expression Omnibus (GEO) repository under accession number GSE212155.

Library generation was performed as described earlier^103, 104^. Briefly, to each sgRNA sequence, BsmBI (Esp3l) cleavage sites sequences (underlined) along with the appropriate BsmBI recognition sequences (CGTCTCA<u>CACC</u>G (sgRNA, 20 nt) <u>GTTT</u>CGAGACG) were added for cloning into the lentiCRISPRv2 (Addgene, #52961) sgRNA and spCas9 expression system. Additional sequences were added for the amplification of the library (italics) and the final oligonucleotide sequence was thus: 5′-(*AGGCACTTGCTCGTACGACG*) CGTCTCA<u>CACC</u>G (sgRNA, 20 nt) <u>GTTT</u>CGAGACG (*ATGTGGGCCCGGCACCTTAA*)-3′. A pool of 5,950 oligonucleotides was synthesized (Twist Biosciences). A primer set annealing to the amplification sequences was used at a final concentration of 0.5 μM to amplify the pool of oligonucleotides using 12.5 μL 2 x NEBnext PCR master mix (New England BioLabs) and 3 μL of oligonucleotide pool (∼3 ng), at a final volume of 25 μL. PCR cycling conditions were 15 s at 98 °C, 20 s at 60 °C, 30 s at 72 °C, for 18 cycles. The resulting amplicons were PCR-purified (NucleoSpin® Extract II, Clontech). To clone the pool of oligonucleotides in the lentiCRISPRv2 vector, we set up Golden Gate assembly reactions at a ratio of 25:1 with 500 ng of vector in reaction buffer (Tango buffer (1x), 0.5 μL DTT (100 mM), 0.5 μL ATP (100 mM), 1 μL (BsmBI) Esp3l (Fermentas), 1 μL T7 ligase) at 50 μL final volume. Golden Assembly reaction was performed using the following cycling conditions: 5 min at 37 °C, 5 min at 20 °C, for 100 cycles. Golden Gate reaction products were isopropanol precipitated using 1 μL of GlycoBlue (15 mg/ml), 4 μL NaCl (5M) and 50 μL isopropanol. After incubation at room temperature for 15 min, we pelleted the library by centrifugation at 13,000 *g* for 20 min, washed the pellet twice with ice-cold 70% ethanol, and centrifuged after each wash at 13,000 *g* for 5 min. After drying, the pellet was resuspended in nuclease-free water. We then performed electroporation using the *E. coli* cells (Endura electrocompetent cells, Lucigen) following the manufacturer’s instructions transforming 90 ng of library DNA into 25 μL of electrocompetent cells for each electroporation reaction for a total of 6 reactions and the cells were pooled and plated on 6 x LB agar plates (245-mm square bioassay dish, 100 μg/ml carbenicillin) and were grown at 30 °C for 20-24 h. Colonies were scraped and plasmid DNA (pDNA) of the sgRNA library was prepared (HiSpeed Plasmid Maxi, Qiagen). Deep-sequencing confirmed that all sgRNAs were well represented in the EpiChromo library. The relative difference in the abundance of sgRNAs in the library at the 10th and 90th percentiles was ∼3.6-fold. sgRNA sequences for the EpiChromo library are presented in Supplementary Table 2.

To generate the virus used for the CRISPR-Cas9 screens, 293FT cells were plated on 150-mm plates and cultured to 80% confluency in order to perform transfection with the pDNA library. Transfection was performed using Lipofectamine 2000 (ThermoFisher) according to the manufacturer’s directions. Briefly, for each transfection in one 150-mm plate, 135 μL of lipofectamine 2000 was added in 4.5 ml of Opti-MEM (Corning) and incubated at room temperature for 5 min. A second solution contained 10.2 μg pMD2.G (Addgene, #12259), 15.6 μg psPAX2 (Addgene, #12260), and 20.4 μg of the library pDNA in 4.5 ml of Opti-MEM. The two solutions were combined and incubated at room temperature for 15-20 min. During this incubation period, the medium on the 293FT cells was changed with 15 ml of fresh prewarmed medium. At the end of the incubation period the transfection mixture was added dropwise to the cells, and then cells were incubated at 37 °C for about 10-12 h after which the transfection medium was removed and replaced with the fresh prewarmed medium. Virus was harvested ∼56 h post-transfection. After clarification of the medium to remove dead cells and debris by centrifugation and passage through 0.45 μm filters, viral particles were collected by using PEG-it Virus precipitation solution following manufacturer’s instructions. Finally, virus was resuspended in mESCs culturing medium and titration was performed using mESCs.

### Establishment and Maintenance CRISPR-Cas9 screens

The screens were performed generally as described previously with some modifications ^103, 104^. Each screen was performed in duplicate. Briefly, for each replicate we used 6 x 10^7^ mESCs carrying the inducible silencing system cultured in medium with 10 μg/ml doxycycline to prevent silencing of the reporter gene. Cells were transduced with the EpiChromo pooled sgRNA library of virus at a multiplicity of infection (MOI) of approximately 0.1 in the presence of polybrene at a final concentration of 5 μg/ml. This results in 6 x 10^6^ transduced mESCs, which is sufficient for the integration of each individual sgRNA into ∼1,000 unique cells. Two days post-transduction, doxycycline (Dox) was removed from the medium to induce silencing of the reporter gene and at the same time 0.25 μg/ml of puromycin was added to the medium to eliminate non-transduced mESCs. The cells were cultured at the minimum number of 6 x 10^6^ cells for each replicate throughout the course of screening in order to ensure coverage of the sgRNAs library in the cell population at ∼1,000 per sgRNA. For the establishment screen, at the end of the establishment period (10 days in -Dox medium), FACS was used to isolate ∼ 3 x 10^6^ eGFP^+^ cells per replicate. Equal numbers of unsorted population of transduced cells were used as controls for comparison. For the maintenance screen, at the end of the establishment period, FACS was used to isolate ∼22 x 10^6^ eGFP^-^ cells per replicate, which were seeded and further cultured. After 16 h of sorting and seeding, the cells were cultured for 3 days in medium with 10 μg/ml Dox to release TetR-KRAB from the 8x-tetO sites and assess how the mutant population of cells perform during the maintenance phase of silencing. The cells were cultured at the minimum number of 6 x 10^6^ cells during the maintenance phase in order to ensure coverage of the sgRNAs library in the cell population at ∼1,000 per sgRNA. At the end of the maintenance period, FACS was used to isolate ∼3 x 10^6^ eGFP^+^ cells per replicate and again equal numbers of unsorted cells were used as controls for comparison. From FACS sorted and unsorted cells for both screens, cell pellets were treated with proteinase K and DNAse-free RNase A in lysis buffer (50 mM KCl, 10 mM Tris-HCl (pH8.3), 2.5 mM MgCl2-6-H2O, 0.45% NP-40, 0.45% Tween-20) and genomic DNA (gDNA) was initially purified using phenol/chloroform/isoamyl alcohol and then ethanol precipitated. PCR of gDNA was performed to attach sequencing adaptors and barcode samples. gDNA of each sample was divided into multiple 50 μL reactions containing a maximum of 2.5 μg gDNA per reaction. For each PCR reaction we used: 0.75 μL of ExTaq DNA polymerase (Clontech), 5 μL of (10x) ExTaq buffer, 4 μL of dNTPs provided with the enzyme, 2.5 μL of P5 stagger primer mix (stock at 10 μM) and 2.5 μL of P7 uniquely barcoded primer (stock at 10 μM). PCR cycling conditions were an initial 3 min at 95 °C; followed by 30 s at 95 °C, 30 s at 53 °C, 30 s at 72 °C, for 23 cycles; and a final 10 min extension at 72 °C. PCR products for each sample were pooled, electrophoresed in an agarose gel, purified (NucleoSpin® Extract II) and sequenced on an Illumina NextSeq 500.

### Screen analysis

The sgRNA counts and abundance were analyzed as described^104^. Guide sequences were extracted from the raw sequencing reads with a modified version of the Python script “count_spacers.py”^104^ using a ‘‘CGAAACACC’’ search prefix, and a counts matrix was generated in each sample. Next, counts files of eGFP^+^ sorted cells and of unsorted population of cells subject to comparison were input into MAGeCK algorithm (version 0.5.9.4) ^38^ and log_2_ fold changes (LFCs) and *P-*values (negative binomial distribution) were calculated for each sgRNA using the ‘mageck test –k’ command with default settings. MAGeCK uses a modified robust ranking aggregation (α-RRA) algorithm to calculate gene level *P*-values and the Benjamini-Hochberg procedure to calculate gene level false discovery rate FDR values^38, 105^. Gene level LFC were calculated by averaging LFCs of all 5 sgRNAs targeting a given gene. The results of this analysis are in Supplementary Tables 3 and 4. The following criteria were used to identify candidate silencing genes with more confidence: i) An FDR value of < 0.1 as a cutoff, ii) at least 3 out of 5 effective (positively selected) sgRNAs, and iii) average of LFCs of the effective sgRNAs (alphamean) higher than 0.585 for the maintenance screen. Using these criteria, we identified 79 and 121 factors affecting silencing during establishment and maintenance, respectively.

Among the 121 factors identified in the maintenance screen, 38 were also found in the establishment screen, and the rest 83 were found to affect silencing only during maintenance based on the above criteria. To increase confidence that we identify maintenance-specific genes, we excluded 19 additional genes (out of 83), which had an effect during establishment at a lower FDR cut-off (FDR < 0.2) than the cut-off originally used (FDR<0.1). Overall, this comparison resulted in the identification of three classes of genes: class I (38 hits, required for both establishment and maintenance), class II (19 hits, maintenance factors with weak effects on establishment), and class III (64 hits, maintenance-specific). Z-scores used in figures were calculated for each sgRNA, Z = (x-m)/s, where x is the LFC for a sgRNA, m is the mean LFC of all sgRNAs, s is the standard deviation of all sgRNAs, and mean Z-score was calculated by averaging the Z-scores of all 5 sgRNAs per gene. CRISPR-Cas9 screen data (fastq and tables) are available at the Gene Expression Omnibus (GEO) repository under accession number GSE212155.

### Functional enrichment and network analysis

Functional enrichment analysis of establishment factors for Figure S2 was performed using g:Profiler (version e106_eg53_p16_65fcd97), with EpiChromo library targeted genes as background list of genes, and Benjamini-Hochberg multiple testing correction method applying significance threshold of 0.05 to retrieve the GO_Biological process (GO_BP) and the GO_Cellular Component (GO_CC) terms enriched^106^.

We performed network analysis using the STRING (version 11.5) database to identify functional interactions (high confidence ≥ 0.85) between the candidate silencing genes^107^. Markov clustering in STRING was performed with an inflation value of 2.5 and array source experiments with no cut off. We used *Mus Musculus* gene symbols and analyzed interactions from all available sources provided by that tool. We performed STRING-based analysis, clustering, and visualization of the network of gene interactions in Cytoscape (version 3.9.0) using a circular layout^108^. Edge weights represent confidence score between gene interaction, and nodes and node weights illustrate genes and log2FC respectively. Colored edges represent clustered gene interactions calculated by Markov clustering. Protein annotations from STRING database were used in Supplementary Tables 3 and 4.

### Screen hit validation

SgRNAs used for the assessment of screen hits were cloned into lentiCRISPRv2 which also encodes the spCas9 gene as described^104^. Individual sgRNAs sequences used to target the indicated genes were the ones used for the pooled CRISRP screens. 4 x 10^4^ mESCs seeded in each well of 96-well plates carrying the inducible silencing system cultured in medium without doxycycline for 6 days prior to infection to induce silencing were individually transduced with lentiviruses carrying one sgRNA per gene in the presence of polybrene at a final concentration of 5 μg/ml. Three days post-infection transduced cells were selected with 1 μg/ml puromycin for three more days in medium without doxycycline. At the end of this selection period, half of the cells for each condition were cultured in the same medium without doxycycline and the other half in the presence of 10 μg/ml doxycycline to assess maintenance. Cells were cultured for three days before assessment of silencing by FACS analysis and the % of eGFP^+^ cells (derepressed cells) was assessed. For the maintenance hit validations, % of derepressed cells was presented over the eGFP^-^ cells shown after establishment. These values were calculated with the formula % = (M^GFP+^ -E^GFP+^) / E^GFP-^ by subtracting GFP^+^ cells during establishment (E^GFP+^) and dividing it with the number of eGFP^-^ cells during establishment (E^GFP-^).

### Flow cytometry

Cells were gently dissociated into single-cell suspension using Accutase (Invitrogen) and resuspended in PBS+1% FBS (FACS medium) and filtered through FACS tubes (Corning™ #352235) with cell strainer cap. FACS was performed using a BD FACSAria II and FACSAria II SORP high speed cell sorters. For silencing assays and screen hit validation, samples were run on a BD FACSCalibur system and the iQue Screener PLUS (IntelliCyt) system. Data were analyzed using FlowJo (version 10.7.2). Single cells were determined by analyzing singlets by comparing cell size (FSC-A) and cell granularity (SSC-A). eGFP gating (eGFP^+^ and eGFP^-^) was determined from the comparison of the distribution of eGFP values from wild-type and *8x-tetO-eGFP* mESCs.

### Generation of knockout mESCs line

To generate *Dhx9* and *Uhrf1* clonal knockout lines in mESCs carrying the inducible heterochromatin system, CRISPR-Cas9 method was applied. sgRNAs were cloned into the lentiCRISPRv2 vector as described^104^ (Supplementary Table 8). Cells were transiently transfected with the plasmid carrying the appropriate sgRNA using Lipofectamine 3000 (ThermoFisher) following manufacturer’s directions. Lipofectamine 3000 and the plasmid were incubated with the cells for 3 h and then replaced with fresh medium. One day after transfection, cells were selected with 1 μg/ml puromycin for three days and then single cells were FACS isolated and seeded in 96-well plates and cultured without puromycin for the rest of clonal expansion. Single clones were expanded and screened by genotyping and MiSeq (Illumina). Knockout clones were validated by western blot.

To generate knockout lines of TetR-3xFlag-KRAB, 2 sgRNAs were used against the *tetR* sequence specifically. sgRNAs were cloned into the lentiCRISPRv2 vector and lentiviruses were prepared. For establishment, cells cultured in doxycycline were transduced simultaneously with the two viruses in the presence of polybrene at a final concentration of 5 μg/ml. Two days post-infection, cells were selected with 1 μg/ml puromycin and cultured in medium without doxycycline to induce silencing. Ten days later, the cells were assessed for silencing by FACS analysis. For maintenance, cells were cultured without doxycycline for 7 days to induce silencing and then were transduced simultaneously with the two viruses in the presence of polybrene. Two days post-infection, selection with puromycin started and one day later doxycycline was added back to the medium. Four days after adding back doxycycline, cells were assessed for maintenance of silencing. Successful knockout was validated with western blot for 3xFlag, and for the maintenance assay was performed two days after puromycin selection (Day 1 of the maintenance assay).

### Protein analysis

Protein samples were loaded on 4-20% gradient TGX Gels (Biorad). SDS-PAGE was performed to separate proteins at 100 Volts for the appropriate amount of time, and proteins were then transferred to a nitrocellulose membrane (Millipore). The membranes were blocked in 5% non-fat dry milk in PBS with 0.2% Tween-20, and sequentially incubated in 1% non-fat dry milk with primary antibodies and HRP-conjugated secondary antibodies, or directly incubated with HRP-conjugated primary antibodies for chemiluminescence detection. The primary antibodies used for western blot analyses were anti-Flag HRP-conjugated (Sigma A8592), anti-GAPDH (Abcam 181603), anti-DHX9 (Abcam 26271), anti-UHRF1 (Santa Cruz sc-373750), anti-TBP (Abcam 818), anti-H3 (Abcam 1791).

### Chromatin immunoprecipitation (ChIP) and quantitative real time PCR (qPCR)

The ChIP assays were performed as described previously^109^. Briefly, to crosslink chromatin, cells were treated with 1% formaldehyde for 10 min at room temperature. Crosslinking was quenched by the addition of glycine to a final concentration of 125 mM for 5 min. The cells were washed twice with ice-cold 1x PBS, scraped into 10 ml of ice-cold PBS, and pelleted by centrifugation at 1000 *g* for 5 min at 4°C. Nuclei were prepared by suspension of cell pellets in a hypotonic buffer (20 mM HEPES pH 7.9, 10 mM KCl, 0.5 mM spermidine, 0.1% Triton X-100, 20% Glycerol, and protease inhibitors), and incubated for 15 min on ice to allow swelling of cells. We then performed dounce homogenization and nuclei were pelleted by centrifugation at 1000 *g* for 5 min at 4°C. The nuclei were lysed in sonication buffer (10 mM Tris-HCl pH 7.5, 1 mM EDTA pH 8.0, 0.1% SDS, and protease inhibitors) and chromatin was sheared with the Covaris E220 Focused-Ultrasonicator (140 watt peak incident, 5% duty factor, 200 cycles/burst, 330 secs sonication time) to a fragment distribution of 200 – 500 bp. Sheared chromatin was diluted with 2x dilution buffer (50 mM HEPES pH 7.9, 280 mM NaCl, 1 mM EDTA, 2% Triton X-100, 0.2% Na-deoxycholate, 0.1% SDS, and protease inhibitors). Chromatin samples were incubated with specific antibodies in the final ChIP buffer (25 mM HEPES pH 7.9, 140 mM NaCl, 1 mM EDTA, 1% Triton X-100, 0.1% Na-deoxycholate, 0.1% SDS, and protease inhibitors) overnight at 4 °C. The protein–DNA complexes were immobilized on pre-washed in ChIP buffer protein A or G dynabeads. For each ChIP, the following antibodies were used with Invitrogen dynabeads: 3 μg of anti-H3K9me3 (Abcam 8898) with protein A, 10 μg of anti-Flag (Sigma F1804) with protein G, 3 μg of anti-H3K27me3 (Millipore, #17622) with protein A, 3 μg of anti-H3K4me3 (Sigma 04-745) with protein A, 6 μg of anti-DHX9 (Abcam 26271) with protein A, 5 μg anti-RNA Pol II (Biolegend (8WG16) MMS-126R-500) with protein A. The bound fractions were washed twice with ChIP buffer, twice with high-salt wash buffer (50 mM HEPES pH 7.9, 500 mM NaCl, 1 mM EDTA, 1% Triton X-100, 0.1% Na-deoxycholate, 0.1% SDS, and protease inhibitors), twice with low-salt wash buffer (20 mM HEPES, 250 mM LiCl, 1 mM EDTA, 0.5% NP-40, 0.5% Na-deoxycholate, and protease inhibitors), and twice with TE buffer (10 mM Tris-HCl pH 8.0, 1 mM EDTA). Elution was carried out with 400 μL elution buffer (50 mM Tris-HCl pH 8.0, 100 mM NaHCO3, 1 mM EDTA, and 1% SDS) at 65 °C for 20 min and then 21 μL of NaCl (4M) was added to each eluate and crosslinks were reversed at 65°C for 10-12 h. After DNase-free RNase A and then proteinase K digestions, DNA samples were purified using phenol/chloroform/isoamyl alcohol and ethanol precipitated with 20 μg glycogen carrier. The precipitated DNA samples were either analyzed by qPCR on Applied Biosystems qPCR instrument in the presence of SYBR green with the primer sequences listed in Supplementary Table 8 using the ΔCT method, or prepared for DNA high throughput sequencing. Reported values are percent of input or occupancy relative to percent input of a negative control region. qPCR quantification and statistical analysis was performed with at least three biological replicates per condition. Data in all figures are presented as mean values +/- S.D. (Standard Deviation).

### H3K9me3 ChIP-seq library preparation and high throughput sequencing

For ChIP-seq, reverse crosslinked DNA treated with RNase A and proteinase K was purified using phenol/chloroform/isoamyl alcohol and ethanol precipitated with 20 μg glycogen carrier. 10 ng of DNA was used to prepare libraries from two biological replicates per sample as described previously^110^. Libraries were pooled and sequenced on Illumina HiSeq platform. Single-end 150-bp raw reads were demultiplexed using the FASTX-Toolkit (version 0.0.13, http://hannonlab.cshl.edu/fastx_toolkit/). Reads were then aligned to the reference mouse genome (GRCm38/mm10, downloaded on January 4, 2020 from https://hgdownload.soe.ucsc.edu/goldenPath/mm10/bigZips/) ^111^ using Bowtie2 (version 2.3.4.3)^112^ in local alignment mode with default parameters. To analyze H3K9me3 signal on the reporter gene, mm10 genome assembly was edited to include 2762 nt of the reporter sequence at its integration site on chromosome 2 (Chr2: 13574347). Chromosomal coordinates in the genome annotation (downloaded on January 13, 2021 from http://hgdownload.soe.ucsc.edu/goldenPath/mm10/bigZips/genes/mm10.ncbiRefSeq.gtf.gz) were edited accordingly to incorporate the reporter.

For H3K9me3 analysis at the reporter gene, duplicate reads were removed using Picard (version 2.8.0) (http://broadinstitute.github.io/picard/). For data visualization, aligned reads were normalized to counts per million using deepTools bamCoverage (version 3.0.2)^113^ and coverage values in bigwig files were computed using 10 bp bins. Genome tracks views were generated in the IGV genome browser^114^. Sequencing data (fastq and bigwig files) are available at the Gene Expression Omnibus (GEO) repository under accession number GSE212155.

### Analysis of differentially methylated H3K9me3 peaks

To analyze differences in H3K9 trimethylation between *Dhx9* KO and wild-type cells, we first defined H3K9me3 peaks/regions using epic2 (version 0.0.51)^62^ for each library with default parameters without removing duplicate reads (-kd) and with wild-type input for background normalization. To minimize biases in peak calling due to varying sequencing depth across libraries, we subsampled aligned reads using SAMtools (version >1.10)^115^ to get ∼61 million reads per library prior to peak calling. Differential read enrichment within H3K9me3 peaks was analyzed using DiffBind (version 3.4.3)^63, 116^ and edgeR. Finally, HOMER annotatePeaks (version 4.11.1)^117^ with default parameters was used to annotate all H3K9me3 peaks.

### Total RNA Purification, reverse transcription (RT), and qPCR

Total RNA from mESCs or NPCs was isolated using TRIzol reagent (Invitrogen, #15596018), treated with TURBO DNase using TURBO DNA-free Kit (Invitrogen, #AM2238) and cleaned up further using RNeasy Mini kit purification (Qiagen) following the manufacturer’s protocol. Total RNA (1 μg) was reverse transcribed using random primers and Superscript III reverse transcriptase (Invitrogen) and quantified by qPCR on Applied Biosystems qPCR instrument in the presence of SYBR green with the primer sequences listed in Supplementary Table 8 using the ΔCT method. *Gapdh* was used as an internal/loading control. Statistical analysis for RT-qPCR was performed on three biological replicates. Data in all figures are presented as mean values +/- S.D.

### Strand specific rRNA-depleted total RNA-seq

Total RNA from mES cells treated with TURBO DNase and cleaned up as described above resulted in high quality RNA (Bioanalyzer RIN > 9.2) that was used for library preparation. rRNA-depleted RNA-seq libraries of 2 biological replicates for wt and *Dhx9* KO mESCs were prepared using TruSeq Stranded Total RNA kit (Illumina) and sequenced on an Illumina HiSeq platform to obtain 150-bp paired-end reads. For differential gene expression analysis, transcript abundances were estimated using kallisto (version 0.45.1)^118^ with strand specificity (--rf-stranded) and bootstrapping (-b 100). Reference sequences of cDNA (fasta format) were obtained from ensembl.org (GRCm38, downloaded in December 2020). Differentially expressed genes in *Dhx9* KO versus wild-type cells were then identified using Sleuth (version 0.30.0)^119^. To visualize alignments on the genome browser, paired-end reads were aligned using HISAT2 (version 2.1.0)^120^ strand specifically (--rna-strandness RF) with default settings. Using SAMtools, high quality (-q10) ‘R1’ or ‘first in pair’ read alignments were extracted (-f64) to prepare bigwig files (deepTools bamCoverage) for visualization in IGV. RNAseq raw data in fastq format and processed bigwig files are available on GEO under accession number GSE212155.

### Analysis of H3K9me3 and RNA expression alterations on repeat elements

To analyze differences in H3K9me3 and RNA expression on repetitive genomic regions, reads were aligned to the mouse genome using Bowtie2 (ChIP-seq) or HISAT2 (RNA-seq) as described above. For RNAseq, only high quality (-q 10) primary alignments (-F 256) of R1 reads (-f 64) were retained (no secondary alignments) using SAMtools. All read alignments (bam files) were converted to bed files using bedtools (version 2.27.1)^121^. Detailed repeat element annotations for the mouse genome were obtained from http://www.repeatmasker.org/species/mm.html (Repeat Library 20140131, downloaded on December 30, 2020). Repeat elements classified as “Simple repeat” or “Low complexity” were excluded from the analysis leaving 3,758,109 annotations comprising 1,362 uniquely named repeat elements (e.g., L1MdTf_I, IAPEY_LTR, MMSAT4, etc.) belonging to 65 repeat families (e.g., LINE/L1, LTR/ERVK, Satellite, etc.).

To analyze changes in H3K9me3 and RNA expression on repeat elements, we compared normalized read counts in wild-type versus *Dhx9* KO cells for both H3K9me3 ChIP-seq and RNAseq. Some repeat elements share high sequence identity (e.g., within the LINE/L1 family, L1MdTf_I, II, and III repeat consensus sequences share nearly 100% sequence identity), which can prove challenging to unambiguously align short reads to individual repeat elements. Considering this limitation, we assigned one alignment per read and summed up the total number of reads per repeat family to estimate methylation enrichment or RNA expression. We then divided normalized read counts per repeat family in *Dhx9* KO by wild-type to get fold differences in expression.

Profile plot for GSAT_MM (major satellite sequence) was prepared by first aligning reads to 471 nt long GSAT_MM consensus sequence (DFAM reference id: DF0003028) ^122^ using Bowtie2 in both wild-type and *Dhx9* KO cells. The alignments were normalized by the total number of reads mapped to the genome (RPM) and converted to bigwig file format using deepTools bamCoverage as described above with a binsize of 1 (-binSize 1) and smoothing the coverage +/- 5 nt of each bin (--smoothLength 5). The profile plots show these alignments/bigwig files as visualized in IGV.

The profile plot for LINE/L1 shows mean read density on all annotated LINE/L1 elements > 6kb length (n = 7023). The plot was created using deepTools (computeMatrix and plotProfile).

### Cellular fractionation and chromatin-associated RNA analysis

Fractionation was performed as previously described with some modifications^123^. Cells from one confluent 15 cm plate (∼2-3 × 10^7^ mES cells) per sample were washed twice with ice-cold 1x PBS and scraped into 10 ml of ice-cold PBS. A fraction of cells (1/10^th^) was kept from each sample to prepare whole cell protein preparations and isolate total RNA. The remaining cells were pelleted by centrifugation at 1000 *g* for 5 min at 4°C. The cell pellets were resuspended in 200 μL of cold lysis buffer (10 mM Tris pH 7.5, 0.15% NP-40, 10 mM KCl) and incubated on ice for 15 min. The lysates were layered onto 500 μL of cold sucrose buffer (10 mM Tris pH 7.5, 150 mM NaCl, 24% sucrose w/v), and centrifuged at 1000 *g* for 10 min at 4°C. The supernatant (cytoplasmic fraction) was not used further. The pellets (nuclei) were resuspended in 200 μL of cold glycerol buffer (20 mM Tris pH 7.9, 75 mM NaCl, 0.5 mM EDTA, 50% glycerol, 0.85 mM DTT) and were gently mixed. Nuclei were lysed by the addition of 200 μL of cold nuclei lysis buffer (20 mM HEPES pH 7.6, 7.5 mM MgCl2, 0.2 mM EDTA, 0.3 M NaCl, 1 M urea, 1% NP-40, 1 mM DTT). The lysates were rotated for 20 min at 4°C and centrifuged at 1000 *g* for 5 min at 4°C. The supernatant from this spin represents the soluble nuclear fraction (nucleoplasm), while the remaining insoluble nuclear fraction represents chromatin. Whole cell preparations, nucleoplasm and chromatin fractions were used for protein analysis with western blotting with the following antibodies were used: anti-DHX9 (Abcam 26271), anti-TBP (Abcam 818), anti-H3 (Abcam 1791).

Whole cell preparations and chromatin fractions of mESCs were used to isolate total and chromatin-associated RNAs, respectively. For RNA isolations we used the TRIzol reagent (Invitrogen, #15596018), and the RNAs were treated with TURBO DNase using TURBO DNA-free Kit (Invitrogen, #AM2238) and cleaned up further using RNeasy Mini kit purification (Qiagen) following the manufacturer’s protocol. Total RNA (300 ng) and chromatin-associated RNAs (200 ng) were reverse transcribed using random primers and Superscript III reverse transcriptase (Invitrogen). Indicated RNAs were quantified by qPCR on Applied Biosystems qPCR instrument in the presence of SYBR green with the primer sequences listed in Supplementary Table 8 using the ΔCT method. *Gapdh* was used as an internal/loading control. Statistical analysis was performed on three biological replicates.

### DNA CpG methylation analysis

Genomic DNA was purified from mESCs and NPCs and 1.5 μg of DNA was bisulfite converted using EpiTect Bisulfite Kit (Qiagen) according to manufacturer’s instructions. PCR products were amplified with EpiTect Methylation-specific PCR (MSP) kit (Qiagen) using the primers shown in Supplementary Table 8, and then subcloned using TOPO Cloning (ThermoFisher) for sequencing. CpG methylation was analyzed with the BISMA tool (http://services.ibc.uni-stuttgart.de/BDPC/BISMA/) using the default parameters.

### mESC growth curves and duplication times

We plated ∼150,000 wild-type or *Dhx9*-KO mESCs in one well of 6-well plate per replicate in the beginning of the time course. At 24, 33, and 48 h after the onset of the course mESCs were collected and stained with trypan blue for counting (Countess^TM^, Invitrogen AMQAX1000). Cell counts were performed for three biological replicates per condition during the time course. Duplication time (DT) was calculated using the following formula: DT = [dT × (ln2)] / [ln (X1 / X0)], (dT: duration (h) between time points (t1: time points) and (t0: time of plating), when the numbers of cells counted are X1 and X0 respectively. For calculating the DT mean values, we used the DT values for all replicates of the three time points counted (n = 9). Data are presented as mean values +/- S.D.

### DNA/RNA hybrid Immunoprecipitation (DRIP)

For DNA/RNA hybrid detection we used mESCs from one confluent 15 cm plate (∼2-3 × 10^7^ mES cells) per sample. Cells were washed twice with ice-cold 1x PBS, scraped into 10 ml ice-cold 1x PBS and pelleted by centrifugation at 1000 *g* for 5 min at 4°C. Nuclei were prepared by suspension of cell pellets in a hypotonic buffer (20 mM HEPES pH 7.9, 10 mM KCl, 0.5 mM spermidine, 0.1% Triton X-100, 20% Glycerol, and protease inhibitors), and incubated for 15 min on ice. We then performed dounce homogenization on ice and nuclei were pelleted by centrifugation at 1000 *g* for 5 min at 4°C. We extracted genomic DNA from the nuclei pellets (∼30-40 μL/sample) by adding 500 μL TE buffer (with 15 μL of 20% SDS, 5 μL proteinase K 20 mg/ml) and incubating overnight at 37°C. Genomic DNA was then isolated with phenol/chloroform/isoamyl alcohol extraction followed by the addition of chilled ethanol and spooling out of DNA. All steps were carried out gently to avoid affecting RNA/DNA hybrids.

Genomic DNA was digested with EcoRI-HF (NEB), XbaI (NEB), BsrGI (NEB), SspI (NEB), Hind III-HF (NEB) overnight at 37°C and the fragmented DNA was purified with phenol/chloroform/isoamyl alcohol. RNA/DNA hybrids were immunoprecipitated from digested DNA (4 μg) using the S9.6 (10 μg) antibody (Kerafast EHN001), or mouse IgG antibody as negative control, and incubated with protein A dynabeads in 1 ml of DRIP buffer (15 mM Tris pH 8.0, 1mM EDTA, 0.01% SDS, 1% Triton, 150mM NaCl). For the RNAse H control IPs, equal amounts of DNA per sample were incubated with RNAse H overnight at 37°C. The bound RNA/DNA hybrids were washed once with DRIP buffer, once with high-salt wash buffer (50 mM HEPES pH 7.9, 500 mM NaCl, 1 mM EDTA, 1% Triton X-100, 0.1% Na-deoxycholate, 0.1% SDS), once with low-salt wash buffer (20 mM HEPES pH 7.9, 250 mM LiCl, 1 mM EDTA, 0.5% NP-40, 0.5% Na-deoxycholate), and once with TE buffer (10 mM Tris-HCl pH 8.0, 1 mM EDTA). Elution was carried out with 300 μL elution buffer (50 mM Tris-HCl pH 8.0, 100 mM NaHCO3, 1 mM EDTA, and 1% SDS) at 65 °C for 20 min and DNA samples were purified using phenol/chloroform/isoamyl alcohol extraction and ethanol precipitation with 20 μg glycogen carrier. The precipitated DNA samples were analyzed by qPCR on Applied Biosystems qPCR instrument in the presence of SYBR green with the primer sequences listed in Supplementary Table 8 using the ΔCT method. Reported values are percent of input. qPCR quantification and statistical analysis was performed with at least three biological replicates per condition. Data are presented as mean values +/- S.D.

## Data availability

CRISRP-Cas9 screen and genome-wide datasets are deposited in the Gene Expression Omnibus (GEO) under the accession number GSE212155.

## Acknowledgments

We thank M. Currie, G. Jih, S. Parhad, T. Shafiq, E. Trompouki, M. Vidaki, and J. Yu for comments on the manuscript, current and former members of the Moazed lab for helpful discussions, and John Doench and Briana Fritchman for providing protocols for CRIPSR library preparation. We thank the Bauer Core Facility at Harvard University for high-throughput sequencing, Jiuchun Zhang of the Initiative for Genome Editing and Neurodegeneration at the Department of Cell Biology at Harvard Medical School for reagents, and the Dana Farber Flow Cytometry core for advice and access to Flow Cytometry equipment. D.M. is an investigator of the Howard Hughes Medical institute.

## Author contributions

A.T. and D.M. conceived the study and designed experiments. A.T. performed all experiments and analyzed the data. H.S analyzed ChIP-seq and RNA-seq data.

A.T. and D.M. wrote the manuscript with feedback from H.S.

---

**Figure S1.**
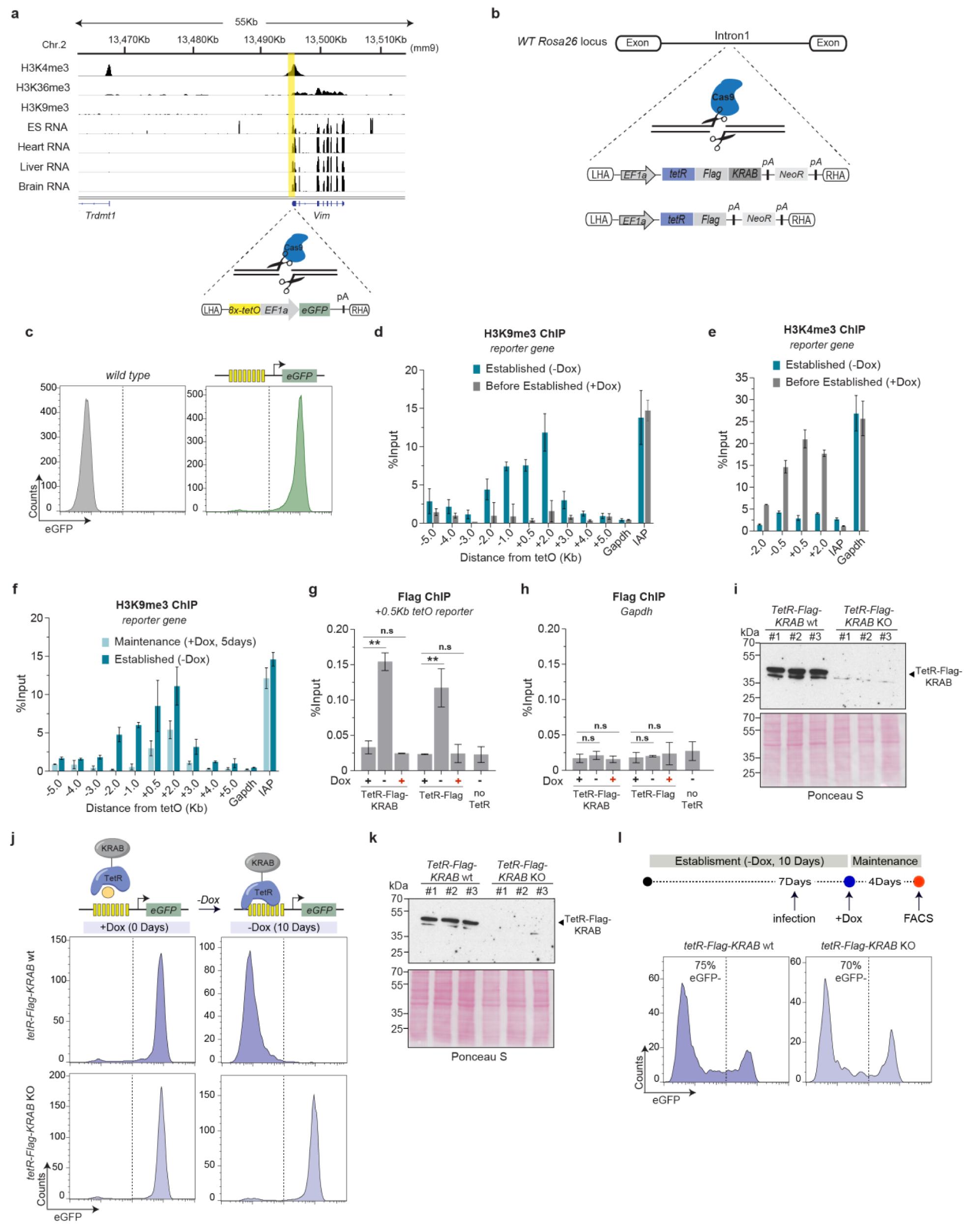
Maintenance of an inducible H3K9me3 domain and silencing of the reporter locus in mESCs. **a,** Histone modifications and RNA expression levels in E14 mESCs and RNA levels in different mouse tissues at the genomic region where *8x-tetO-eGFP* reporter gene was inserted located on chromosome 2. Data from the ENCODE project are displayed and the genome coordinates are based on mm9 (MGSCv37) genome assembly. The *8x-tetO-eGFP* reporter gene was integrated via HDR-mediated CRISPR-Cas9 at the promoter of the vimentin gene (Vim) as depicted below the tracks. LHA and RHA, left and right homology arms; pA, transcription termination signal. **b,** Schematic diagram depicting the strategy and integration site of TetR-Flag-KRAB and TetR-Flag in the first intron at the *Rosa26* locus located on chromosome 6 in mESCs. **c,** Flow cytometry histograms show eGFP expression in wild-type and *8x-tetO-eGFP* mESCs in the absence of TetR fusion proteins. **d,** ChIP-qPCR analysis for H3K9me3 at the *8x-tetO-eGFP* reporter locus and surrounding regions in mESCs expressing TetR-Flag-KRAB cultured in the presence (Before Established, +Dox) or absence (Established, -Dox) of doxycycline. *Gapdh* and *IAP*, are used as controls for euchromatin and heterochromatin H3K9me3 levels respectively. Values are shown as percentage (%) of input. Error bars, standard deviation (SD); n = 3 replicates. **e,** Same as in **d** for H3K4me3 ChIP. **f,** Same as in **d** for H3K9me3 levels in mESCs cultured in the absence (Established, -Dox) or presence for 5 days of doxycycline (Maintenance, +Dox). **g,** ChIP-qPCR analysis for TetR-Flag-KRAB and TetR-Flag binding at the *8x-tetO-eGFP* reporter locus before establishment (+Dox), after establishment (-Dox), and during the maintenance phase of silencing after adding back doxycycline to the medium for 24 hours (+Dox, in red). *8x-tetO-eGFP* reporter mESC line without TetR-Flag-KRAB or TetR-Flag was used as a negative control. Values are shown as percentage (%) of input. Error bars, standard deviation (SD); n = 3 biological replicates. **h,** Same as in **g** at the *Gapdh* promoter. **i,** Western blot (top) showing protein levels of TetR-Flag-KRAB before (*TetR-Flag-KRAB* wt) and after its deletion (*TetR-Flag-KRAB* KO) in mESCs cultured in the presence of doxycycline before establishment of silencing. Ponceau S staining used as a loading control (bottom). Molecular weights in kilodalton are shown on the left. **j,** Flow cytometry histograms showing eGFP expression in *tetR-Flag-KRAB* wt and *tetR-Flag-KRAB* KO mESCs cultured with and without doxycycline (-Dox). **k,** Same as in **i** but TetR-Flag-KRAB was deleted after establishment of silencing. **l,** Same as in **j** showing eGFP expression in mESCs with the indicated genotypes. Top, diagram of experimental strategy. Cells were cultured for seven days without doxycycline (-Dox), followed with infection with two lentiviral vectors carrying two sgRNAs targeting the TetR domain of TetR-Flag-KRAB, three days later doxycycline (+Dox) was added back for four days. Bottom, Flow cytometry histograms showing similar maintenance of silencing in in *tetR-Flag-KRAB* wt and *tetR-Flag-KRAB* KO mESCs.

**Figure S2.**
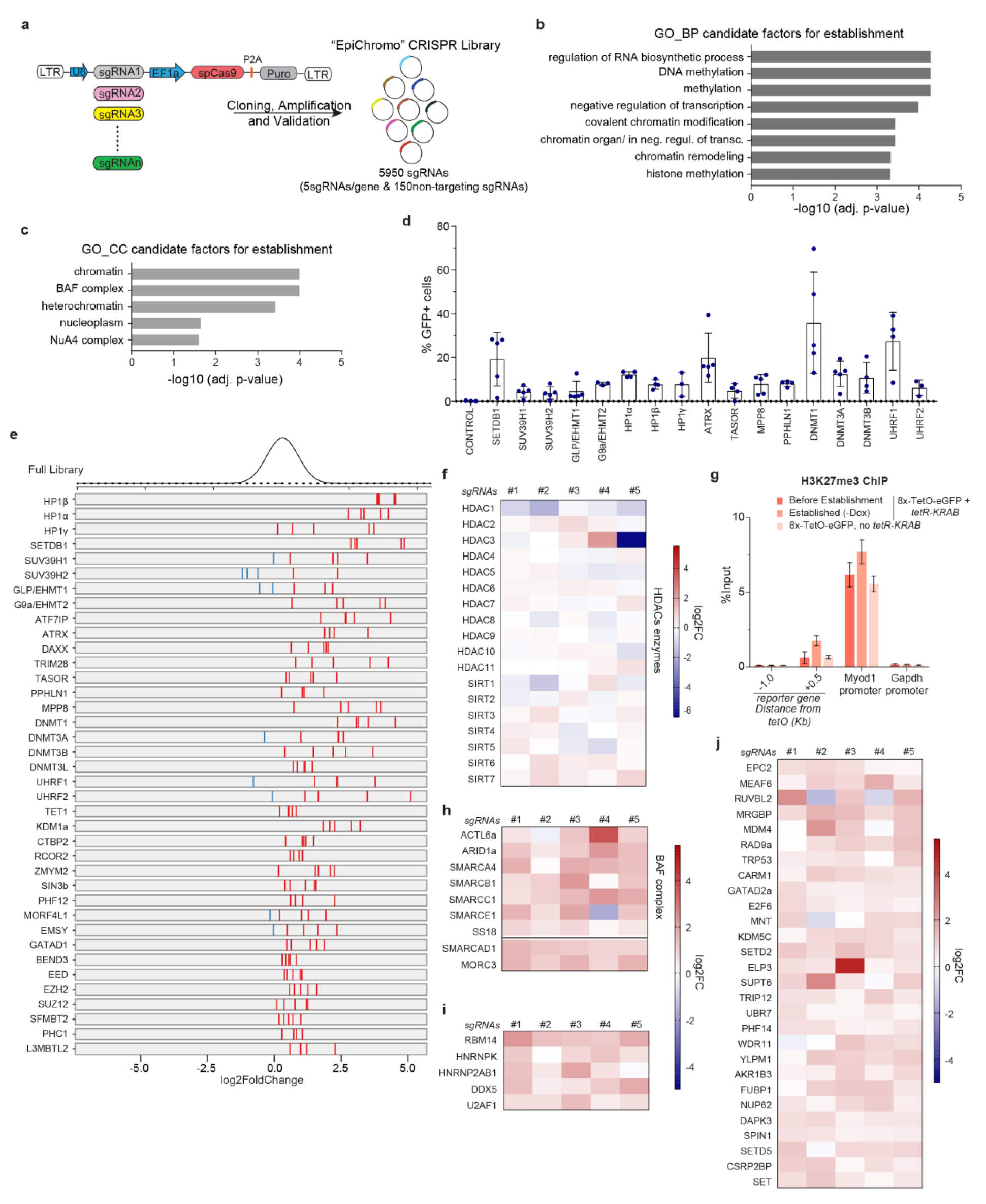
Supplemental of establishment screen data. **a,** Schematic diagram showing the targeting vector and strategy followed to generate the EpiChromo library used to perform the pooled screens. **b,** Gene ontology (GO_BP) analysis for the biological processes associated with the 79 establishment factors. **c,** Gene ontology (GO_CC) analysis for the cellular components associated with the 79 establishment factors. Of note, the NuA4 complex shares components with HDAC complexes likely explaining its appearance on the list of candidate factors. **d,** Validation of candidate genes from the screen associated with H3K9 methylation and DNA methylation. The indicated genes were targeted by three to five sgRNAs individually and eGFP expression was measured by FACS. Percentage (%) of eGFP^+^ cells are presented. Mean values are shown, error bars are SD; n=3-5 sgRNAs each gene. **e,** Performance in the pooled screen of sgRNAs targeting the 38 genes depicted in Figure 2e. Log_2_ fold change in eGFP^+^ sorted versus unsorted cells is shown for the full library of sgRNAs (top) and for the 5 sgRNAs targeting each gene. sgRNAs with positive or negative log_2_ fold change are shown in red or blue respectively. **f,** Heatmap depicting HDAC enzymes (left) affecting establishment of silencing. Log2 fold change of each one of the 5 sgRNAs targeting each gene in eGFP^+^ sorted versus unsorted cells is shown (n = two independent replicates). **g,** ChIP-qPCR analysis for H3K27me3 at the *8x-tetO-eGFP* reporter locus in mESCs before establishment, after establishment of silencing and in *8x-tetO-eGFP* mESCs without TetR-Flag-KRAB. Values are shown as percentage (%) of input. *Myod1* and *Gapdh* are used as H3K27me3 positive and negative control regions. Error bars, standard deviation (SD); n = 3 biological replicates. **(h-j)** Same as in **f** for chromatin remodelers, RNA processing factors and cluster 8 genes respectively.

**Figure S3.**
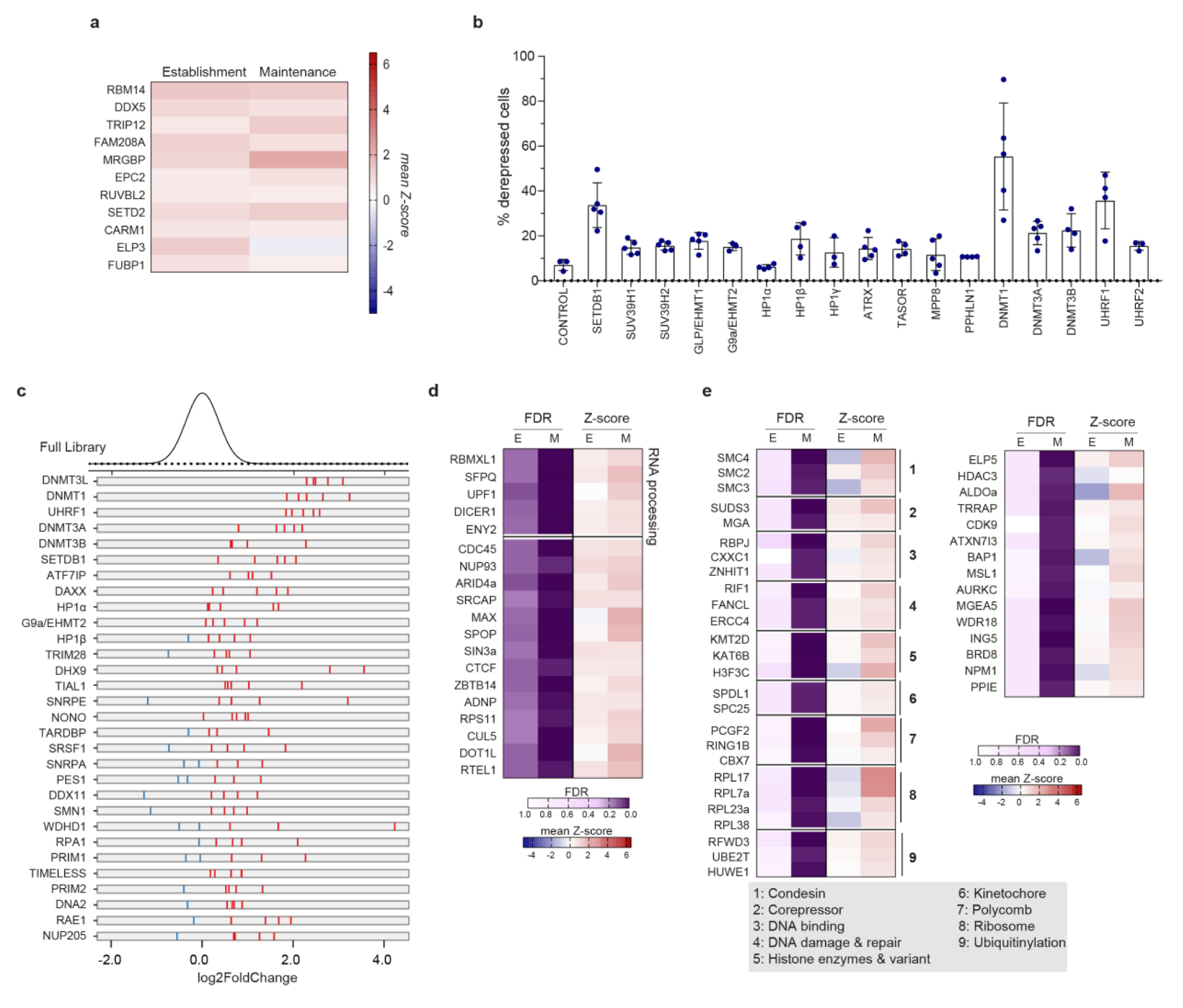
Analysis of maintenance screen data. **a,** Heatmap of 15 out of the 38 gene hits (left) that affect both establishment and maintenance of silencing and are not presented in Figure 3d. Mean values of the Z-scores of all 5 sgRNAs targeting each gene in eGFP^+^ sorted versus unsorted cells are shown in each condition (n = two independent replicates). **b,** Validation of candidate genes from the screen associated with H3K9 and DNA methylation. The indicated genes were targeted by three to five sgRNAs individually and eGFP expression was assessed by FACS. Percentage (%) of eGFP^+^ cells is indicated (y axis). Mean values are shown, error bars are SD; n=3-5 sgRNAs each gene. **c,** Performance in the pooled screen of sgRNAs targeting the gene hits shown (left). Log_2_ fold change in eGFP^+^ sorted versus unsorted cells is shown for the full library of sgRNAs (top) and for the 5 sgRNAs targeting each gene. sgRNAs with positive or negative log_2_ fold change are shown in red or blue respectively. **d,** Heatmap of the 19 gene hits found to affect both maintenance (FDR < 0.1) and establishment but at a lower cutoff (FDR < 0.2). Gene-level FDR values and mean values of the Z-scores of all 5 sgRNAs targeting each gene in eGFP^+^ sorted versus unsorted cells during establishment (E) or maintenance (M) are shown. (n = two independent replicates). **e,** Same as in **d** showing gene hits (left) belonging to the indicated functional categories (right) that are maintenance-specific regulators of silencing.

**Figure S4.**
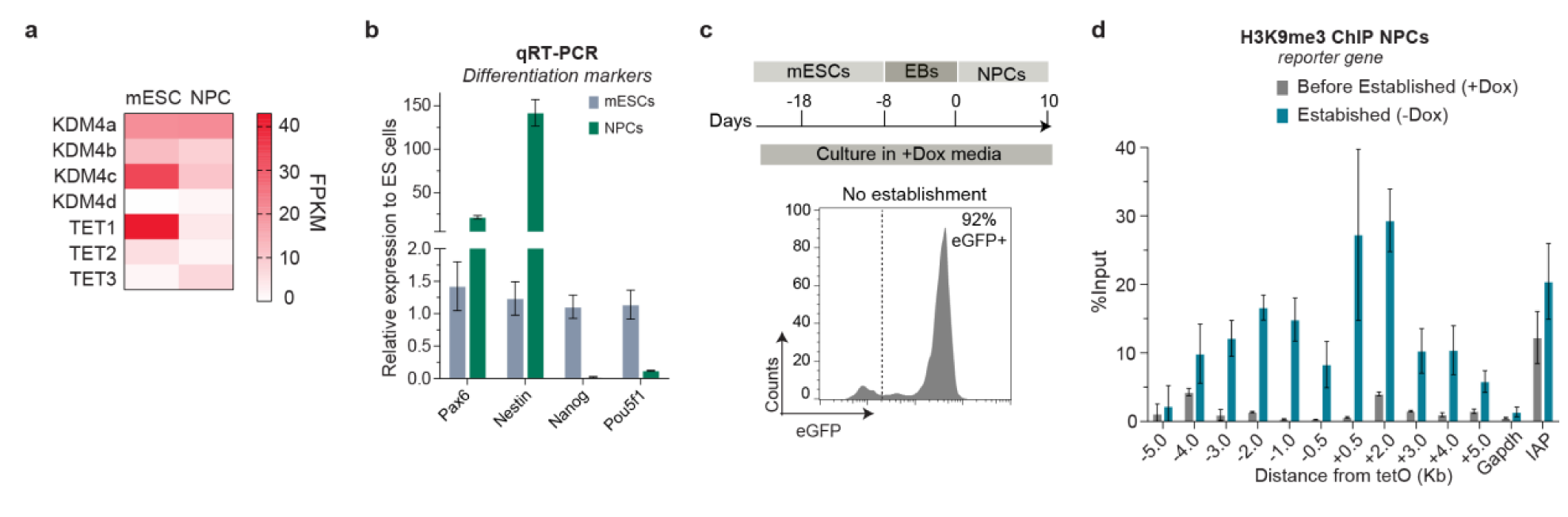
Epigenetic maintenance of silencing is stabilized upon differentiation. **a,** Heatmap showing expression levels of KDM4s and TETs in mESCs and NPCs quantified by RNA-seq in^56^. Values are displayed as fragments per kilobase of transcript per million fragments mapped (FPKM). **b,** qRT-PCR analysis showing increased expression levels of neural progenitor cell markers (*nestin, pax6*) and decreased levels of embryonic stem cell related genes (*nanog, pou5f1*) after differentiation of *8x-tetO-eGFP/TetR-Flag-KRAB* reporter mESCs to NPCs. Mean values are shown. Error bars, standard deviation (SD); n = 3 biological replicates. **c,** Flow cytometry histogram show eGFP expression in neural progenitor cells that were differentiated from mESCs in the continuous presence of doxycycline (+Dox). Percentage (%) indicates fraction of eGFP^+^ cells. **d,** ChIP-qPCR analysis for H3K9me3 at the *8x-tetO-eGFP* reporter locus and surrounding regions in neural progenitor cells (NPCs) cultured in the presence (Before Established, +Dox) or absence (Established, -Dox) of doxycycline. *Gapdh* and *IAP*, are used as controls for euchromatin and heterochromatin H3K9me3 levels respectively. Values are shown as percentage (%) of input. Error bars, standard deviation (SD); n = 3 replicates.

**Figure S5.**
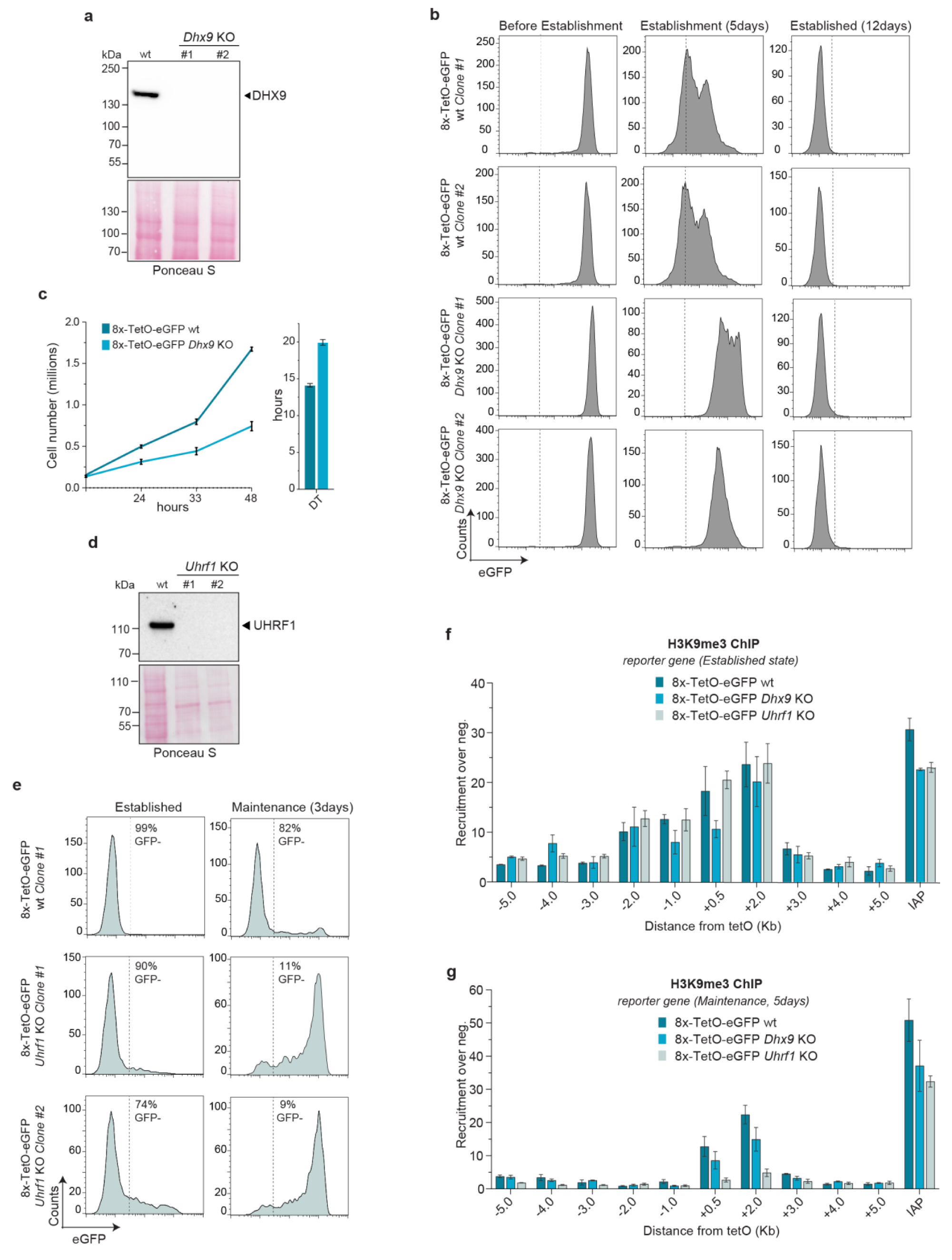
Deletion of *Dhx9* results in slower establishment and defective maintenance of H3K9me3. **a,** Western blot showing (top) protein levels of DHX9 in *8x-tetO-eGFP* wild-type (wt) and two different clones (#1, #2) of *Dhx9* KO mESCs and (bottom) Ponceau S staining used as a loading control. Molecular weights in kilodalton are shown on the left. **b,** Flow cytometry histograms show eGFP expression in *8x-tetO-eGFP* wild-type (wt) and *Dhx9* KO mESCs before and after culturing cells in medium with doxycycline for 5 and 12 days. **c,** Left, growth curves of *8x-tetO-eGFP* wild-type (wt) and *Dhx9* KO mESCs. Right, duplication time (DT) of the indicated genotypes calculated based on the curve data. Mean values are shown; error bars, SD. **d,** Western blot showing (top) protein levels of UHRF1 in *8x-tetO-eGFP* wild-type (wt) and two different clones (#1, #2) of *Uhrf1* KO mESCs and Ponceau S staining used as a loading control (bottom). Molecular weights in kilodalton are shown on the left. **e,** Flow cytometry histograms showing eGFP expression in *8x-tetO-eGFP* wild-type (wt) and *Uhrf1* KO mESCs cultured in medium without doxycycline for 12 days (Established State) and three days after adding back doxycycline during the maintenance phase. Percentages (%) indicate fraction of eGFP^-^ cells. **f,** ChIP-qPCR analysis for H3K9me3 at the *8x-tetO-eGFP* reporter locus and surrounding regions in *8x-tetO-eGFP* wild-type (wt), *Dhx9* KO, and *Uhrf1* KO mESCs cultured in the absence (Established state, -Dox) of doxycycline. Values are shown relative to *Gapdh* used as a negative control region. Error bars, standard deviation (SD); n = 3 replicates. **g,** Same as in **e** but cells are cultured in the presence of doxycycline (+Dox) for 5 days during the maintenance phase.

**Figure S6.**
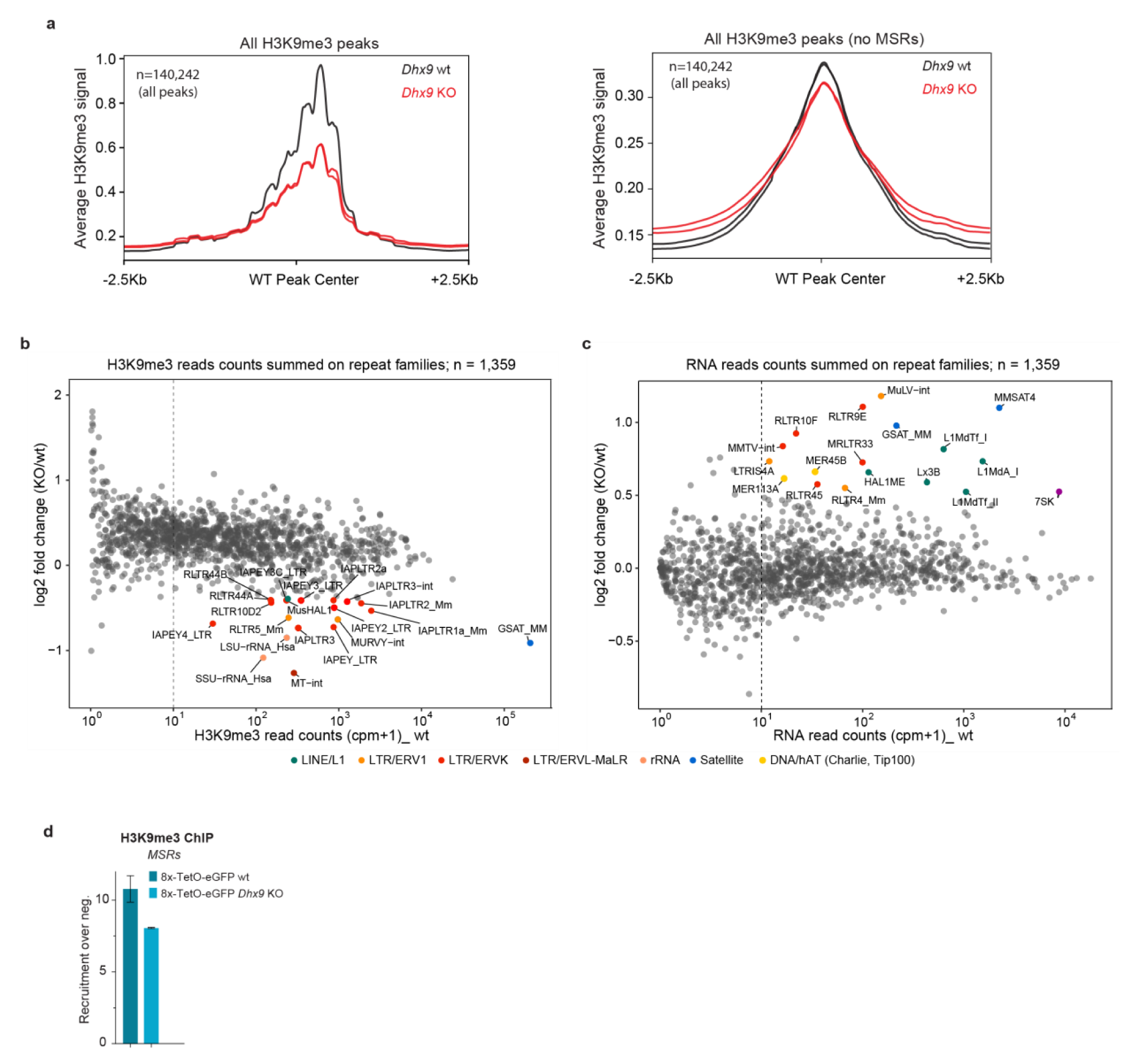
Effects of deleting DHX9 on endogenous heterochromatic regions. **a,** Plot showing the normalized average density of H3K9me3 reads across all H3K9me3 peaks in wild-type and *Dhx9* KO samples (left). Plot showing the normalized average density of H3K9me3 reads across all H3K9me3 peaks excluding MSRs (right). **b,** Scatterplot showing mean H3K9me3 read counts in wt mESCs (x-axis) for 1,359 repeat elements and log_2_ ratio of total H3K9me3 ChIP-seq reads (y-axis) per repeat element in *Dhx9* KO (KO) versus wild-type (wt) mESCs. Names of selected repeat elements are denoted, and respective repeat family is color-coded. Source data for this figure are provided in Supplementary Table 6. **c,** Scatterplot showing mean RNA read counts in wt mESCs (x-axis) for the 1,359 repeat elements plotted against log_2_ ratio of total RNA-seq reads (y-axis) per repeat element in *Dhx9* KO (KO) versus wild-type (wt) mESCs. Names of selected repeat elements are denoted, and respective repeat family is color-coded. Source data for this figure are provided in Supplementary Table 6. **d,** ChIP-qPCR analysis for H3K9me3 at major satellite repeats in *8x-tetO-eGFP* wild-type (wt) and *Dhx9* KO mESCs. Values are shown relative to *Gapdh* used as a negative control region. Error bars, standard deviation (SD); n = 3 replicates.

**Figure S7.**
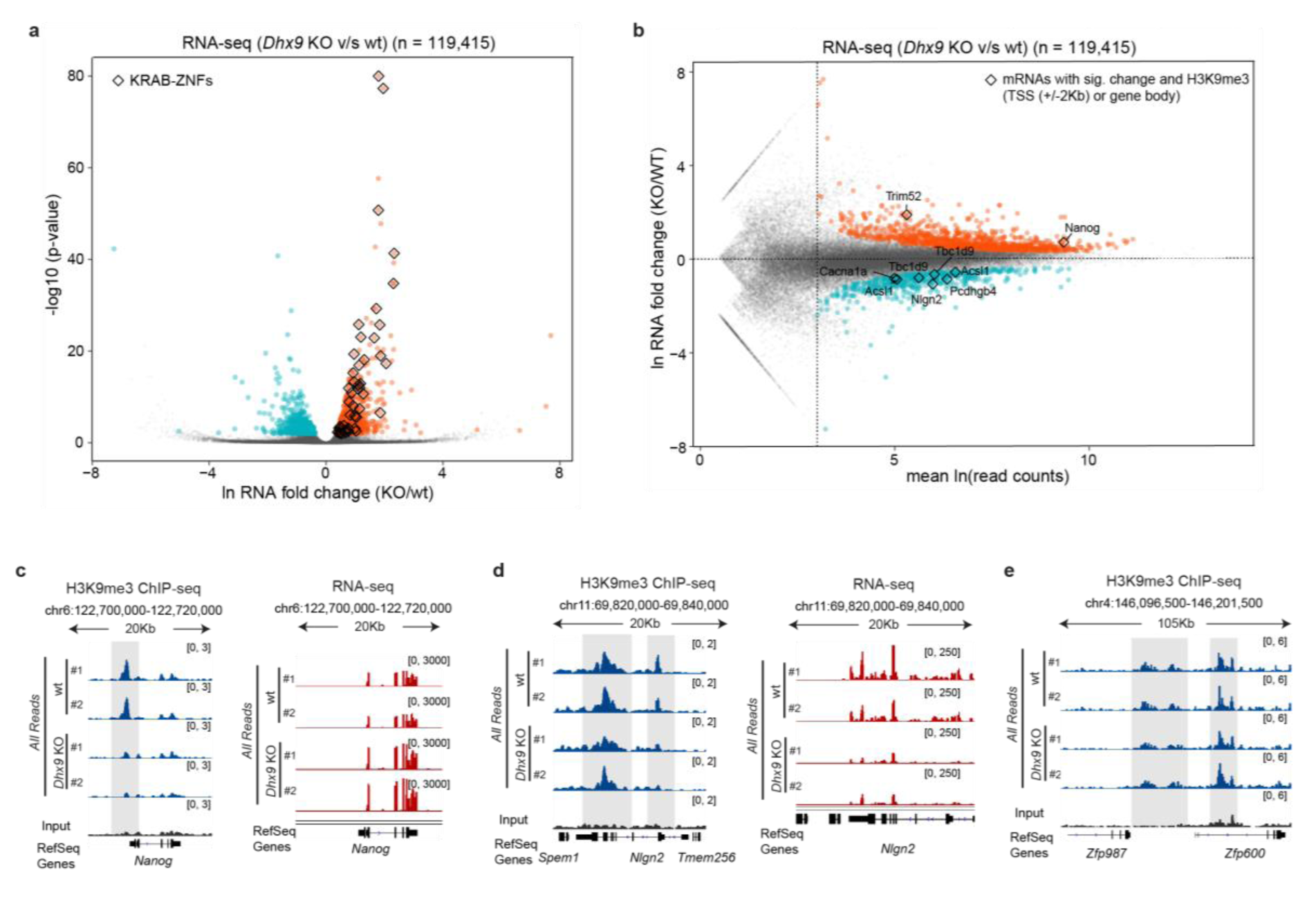
Effects of deleting DHX9 on endogenous heterochromatic regions. **a,** Volcano plot displaying the log (ln) fold difference in transcript abundance in *Dhx9* KO (KO) versus wild-type (wt) mESCs (x-axis) plotted against the negative log_10_ p-value (y-axis) for 119,415 annotated RNA transcript isoforms. Using a q-value cut-off of < 0.01, 1018 transcripts show increased expression (orange) and 506 are decreased (blue) in *Dhx9* KO relative to wild-type mESCs. KRAB-ZNFs genes are highlighted with diamonds. **b,** Scatter plot comparing log (ln) mean RNA read counts in all samples for 119,415 annotated RNA transcript isoforms (x-axis) and the log (ln) fold difference in transcript abundance in *Dhx9* KO (KO) versus wild-type (wt) mESCs (y-axis). Using a q-value cut-off of < 0.01. Genes with a significant change of their RNA levels that overlap with H3K9me3 around their promoter regions (+/-2 kb TSS), or their gene body are highlighted with diamonds. Source data for this figure are provided in Supplementary Table 7. (**c** and **d**) Genome tracks of ChIP-seq for H3K9me3 (left) and total RNA-seq (right) at the indicated locus in *8x-tetO-eGFP* wild-type (wt) and *Dhx9* KO mESCs. Normalized reads are presented in brackets. All reads, uniquely mapped reads plus one alignment reported for reads mapping to multiple genomic locations. Top, chromosome coordinates. **e,** Genome tracks of ChIP-seq for H3K9me3 at the indicated locus in *8x-tetO-eGFP* wild-type (wt) and *Dhx9* KO mESCs. Normalized reads are presented in brackets. All reads, uniquely mapped reads plus one alignment reported for reads mapping to multiple genomic locations. Top, chromosome coordinates.

**Figure S8.**
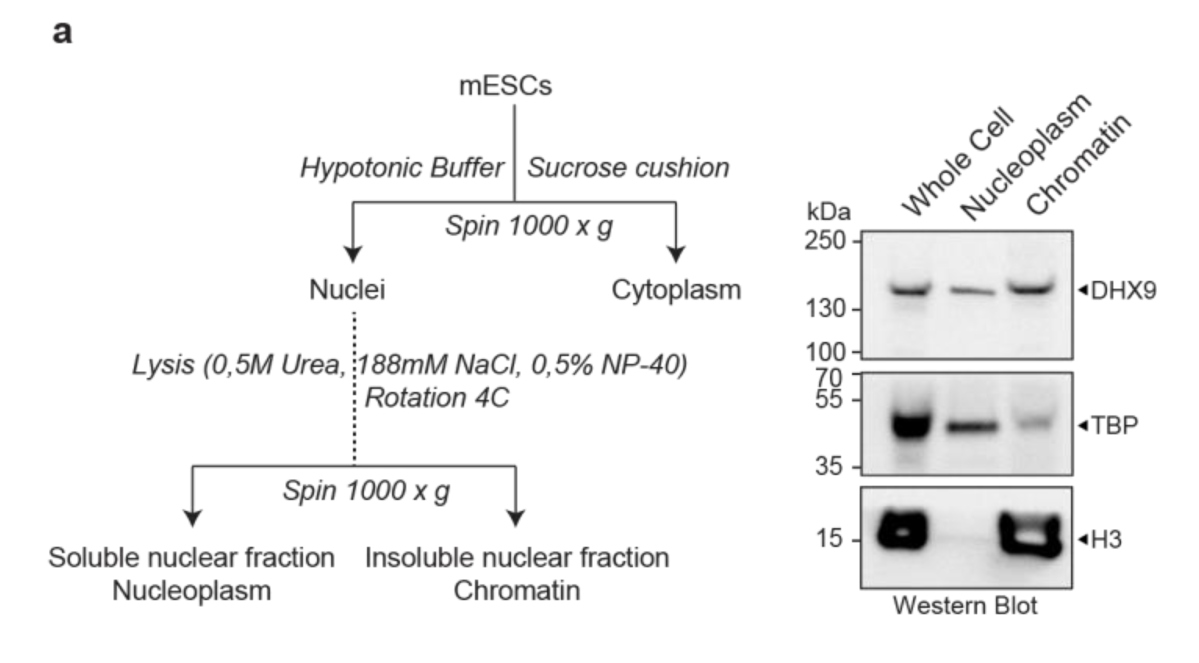
The DHX9 protein is enriched in the chromatin fraction. **a,** Left, schematic of the strategy used for the biochemical separation of the chromatin fraction used for the isolation of RNA and protein. Right, western blot showing the levels of DHX9 (top), TBP (middle), and H3 (bottom) proteins in the indicated cellular fractions. Molecular weights in kilodalton are shown on the left.

